# Elongin B orchestrates chromatin and transcriptional programs in H3K27M-mutant diffuse midline glioma

**DOI:** 10.64898/2026.05.16.724973

**Authors:** Alan L. Jiao, Rebecca L. Murdaugh, Adam F. Kebede, Claudia A. Mimoso, Barry M. Zee, Rosemary U. Richard, Kwanha Yu, Caitlin L. Bagnetto, Karen Adelman, Mariella G. Filbin, Yang Shi, Jamie N. Anastas

**Author notes:** denote current affiliations. equal contribution (Alan L. Jiao, Rebecca L. Murdaugh, Adam F. Kebede).

## Abstract

Recurrent driver mutations in genes encoding histone H3 (H3.3K27M and H3.1K27M) are observed in ∼80% of diffuse midline gliomas (DMG), which lead to aberrant gene regulation, yet the specific RNA polymerase II (Pol2) regulators that induce transcriptional dysregulation in DMG are not fully defined. We identified multiple regulators of Pol2 elongation as DMG genetic dependencies in a chromatin-focused CRISPR screen. Additional studies confirm that knockout (KO) of the Pol2 SIII complex gene elongin B (ELOB) inhibits DMG cell proliferation in tissue culture and tumor growth in xenograft models. Further genomic analyses reveal that ELOB binding sites are enriched in H3K27M oncohistones and that ELOB KO alters H3K27me3 and H3K27M incorporation at thousands of genomic regions, implicating ELOB in the maintenance of dysfunctional chromatin states in DMG. Correspondingly, PRO-seq and RNA-seq profiling reveal that ELOB loss disrupts Pol2 transcriptional activity and alters the expression of transcripts involved in metabolism, proliferation, and brain development. These findings suggest that Pol2 elongation factors like ELOB cooperate with H3K27M oncohistones to maintain the epigenetic and transcriptional landscape driving DMG malignancy.

## INTRODUCTION

H3K27M mutations are present in ∼80% of diffuse midline glioma (DMG), a group of highly aggressive brain tumors diagnosed in young children with no effective therapies (*1*, *2*). H3K27-altered DMG are characterized by disruption of Polycomb-dependent gene silencing through H3K27me3 and a gain of histone acetylation associated with gene activation, leading to aberrant transcriptional programs that drive tumorigenesis (*3–6*). Malignant gliomas are dependent on atypical RNA polymerase II (Pol2) transcriptional activity, as highlighted in recent studies establishing roles for Pol2 elongation regulators like CDK9 and AFF4 in sustaining aberrant gene regulatory programs in DMG and other brain tumors (*7–11*). How specific Pol2 regulators cooperate with H3K27M oncohistones and other epigenetic and genetic drivers to enforce aberrant transcription in DMG remains poorly understood.

Dynamic regulation of RNA polymerase II (Pol2) activity is integral to the execution of gene expression programs in both normal cells and cancer and can occur at multiple stages of the transcription cycle, including during transcriptional initiation and elongation. Pol2 elongation is regulated in part by specialized complexes such as the super elongation complex (SEC) and the Elongin (SIII) complex (*12*, *13*). Due to their ability to enhance Pol2 transcription in enzymatic assays performed using reconstituted mixtures of purified chromatin, enzymes, and cofactors (*13–18*), it was long assumed that many Pol2 elongation regulators would act as general transcriptional factors controlling the expression of large numbers of target genes. However, recent studies evaluating Pol2 elongation factor function in different tissue contexts call this assumption into question, as genetic inhibition of Pol2 regulators, like *ELOA* and *ELOB,* does not change Pol2 elongation rate on a global scale in mammalian cells but rather affects the expression of select transcripts, such as genes involved in stem cell biology and cell fate specification (*19–23*). These findings suggest that certain Pol2 regulators might be hijacked to activate cancer-specific gene expression programs.

We recently performed chromatin- and transcription-focused CRISPR dropout screens (*24*) that identified multiple Pol2 elongation regulators, including *Elongin B* (*ELOB*), as DMG genetic dependencies. Follow-up studies confirm that ELOB promotes DMG tumor growth *in vitro* and in an orthotopic xenograft model. Further chromatin profiling, PRO-seq, and RNA-seq studies demonstrate a role for ELOB in maintaining DMG disease-associated chromatin and gene expression programs involved in metabolic regulation, mitosis, and neural and glial differentiation.

## RESULTS

### The Pol2 SIII complex is a genetic dependency in H3K27M-mutant diffuse midline glioma (DMG)

To identify genes driving the growth and survival of DMG, we conducted chromatin-focused CRISPR dropout screens in H3K27M-mutant SU-DIPGVI and SU-DIPGXIII cells. Stable Cas9-expressing SU-DIPGVI and SU-DIPGXIII cells were infected with a pooled sgRNA library targeting 1,354 chromatin readers, writers, erasers, remodelers, and other epigenetic and transcriptional regulators. Analysis of sgRNA abundance identified a consensus of 149 sgRNA “hits” that significantly inhibited the growth of both SU-DIPGXIII and SU-DIPGVI cell lines (\**p*<0.01, **fig. S1, A** and **B**) (*24*). We used the STRING protein-protein interaction database and gene ontology (GO) analysis to cluster top screen hits according to annotated functional categories and associations with known protein complexes. Genes involved in the RAN (RAS-related nuclear protein) pathway, mRNA splicing, histone acetylation, and RNA Polymerase II (Pol2) elongation were observed among the top DMG genetic dependencies (**Fig. 1A** and **fig****. S1, C** and **D**). Members of the Pol2 SIII elongation complex, which functions to promote Pol2 transcription (**Fig. 1B**) (*15*, *25*), including *Elongin B* (*ELOB*) (SU-DIPGVI: *\*\*\*\*p*<1.665×10^−5^, SU-DIPGXIII: \*\*\**p*<0.000101) and its binding partner *Elongin C* (*ELOC*) (SU-DIPGVI: \**p*<0.026, SU-DIPGXIII: \**p*<0.030), were identified among the strongest DMG genetic dependencies and selected for further study (**fig. S1E**).

**Figure 1:**
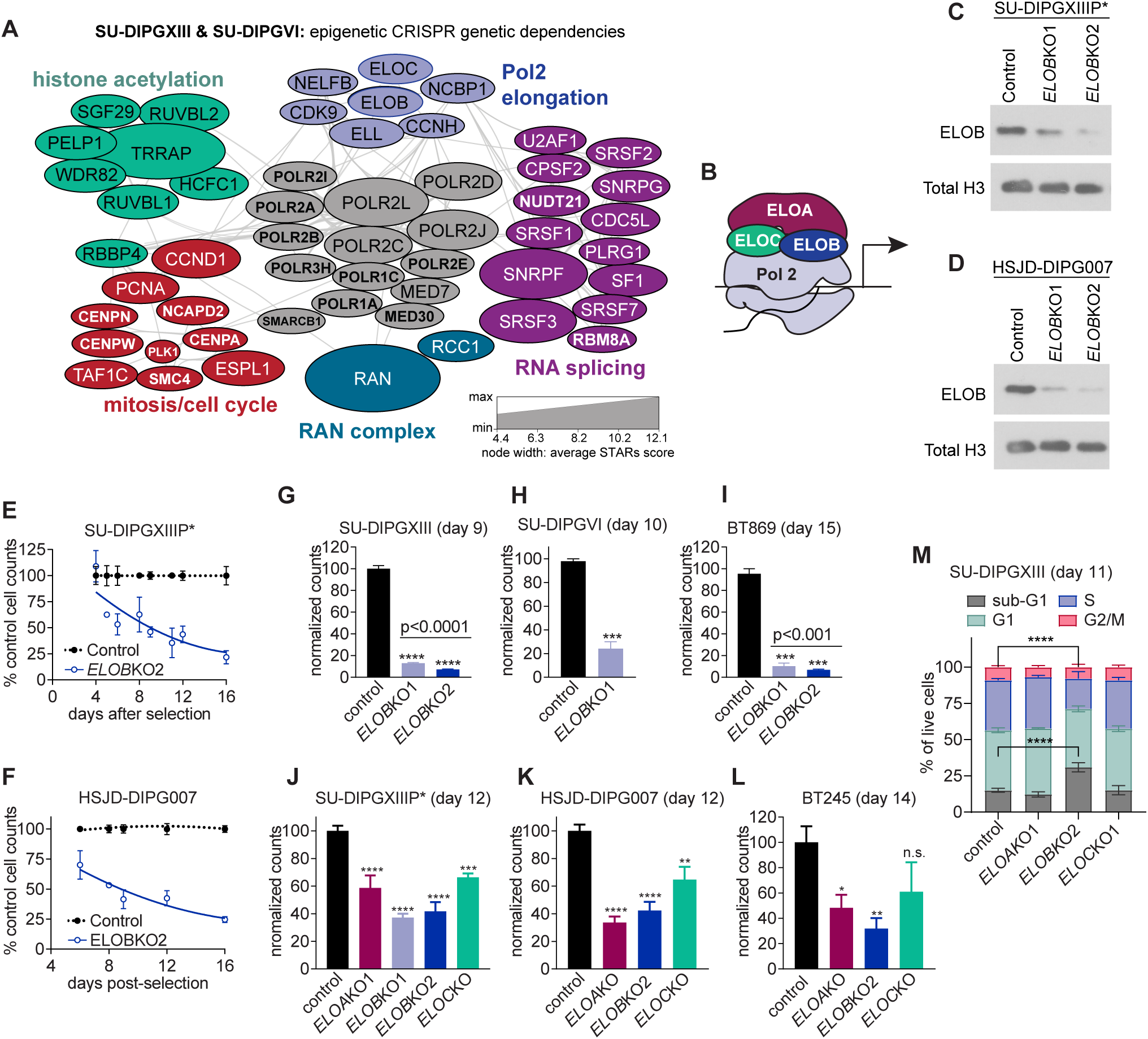
The Pol2 SIII complex is a genetic dependency in H3K27M-mutant diffuse midline glioma (DMG). **(A)** Protein-protein interaction network illustrating selected genetic dependencies from a chromatin-focused CRISPR dropout screen in two H3K27M-mutant glioma cell lines (SU-DIPGXIII and SU-DIPGVI) highlighting roles for regulators of Pol2 elongation, histone acetylation, and RNA splicing in DMG growth and survival. Node width is scaled to the average STARS score, with larger sizes correlating with stronger genetic dependencies. **(B)** Schematic of RNA polymerase II (Pol2) SIII transcriptional regulatory complex of ELOA, ELOB, and ELOC. **(C** and **D)** Immunoblot analysis revealing reduced ELOB protein in SU-DIPGXIIIP*+Cas9 **(C)** and HSJD-DIPG007+Cas9 DMG **(D)** cells transduced with two independent *ELOB* sgRNAs (*ELOB*KO1 and KO2). **(E** and **F)** Normalized cell counts of control and *ELOB*KO SU-DIPGXIIIP* **(E)** and HSJD-DIPG007 **(F)** cells at various time points following selection. **(G** to **I)** Normalized cell counts in Cas9 (+) H3K27M-mutant SU-DIPGXIII and SU-DIPGVI cells used in the original screen and H3K27M mutant BT869 cells, expressing either control or *ELOB* sgRNAs. **(J** to **L)** Normalized cell counts of Cas9 (+) H3K27M-mutant DMG cell lines following CRISPR-based targeting of *ELOA*, *ELOB,* or *ELOC*. **(M)** BrdU pulse labeling and cell cycle analysis showing the average percent of live SU-DIPGXIII+Cas9 cells transduced with control, *ELOA*, *ELOB*, or *ELOC* sgRNAs in different cell cycle phases, suggesting reduced BrdU(+) S-phase cells and increased sub-G1 cells following *ELOB* loss. Error bars show S.E.M. with *p<0.05, **p<0.01, ***p<0.001, and ****p<0.0001 as determined by one-way ANOVA or Student’s t-test.

### The ELOA/ELOB/ELOC complex promotes DMG cell growth

To validate the findings of our initial CRISPR screen, we evaluated the impact of *ELOB* knockout (*ELOB*KO) across multiple additional patient-derived H3K27M-mutant DMG cell lines maintained in serum-free neurosphere culture. We used lentiviral transduction to introduce two independent sgRNAs (*ELOB*KO1 and *ELOB*KO2) into Cas9(+) SU-DIPGXIIIP*, an aggressive DMG model derived from an invasive SU-DIPGXIII subpopulation (*26*), and Cas9(+) HSJD-DIPG007 DMG cells, which harbor *ACVR1* and *PPM1D* mutations in addition to the H3K27M alteration. Western blotting and immunofluorescence microscopy confirmed a 60-90% reduction in ELOB in the *ELOB*KO cells (**Fig. 1, C** and **D**, and **fig. S1F**). Longitudinal growth analyses of these polyclonal *ELOB*KO cells revealed a progressive loss of the *ELOB*KO cells in both the SU-DIPGXIIIP* and HSJD-DIPG007 cell line models compared to controls (**Fig. 1, E** and **F**). These validated *ELOB* sgRNAs also led to a 75–90% reduction in growth of the SU-DIPGXIII and SU-DIPGVI cell lines used in the original screen and similarly inhibited the proliferation of H3K27M-mutant BT869 cells (**Fig. 1, G** to **I**). These results align with prior studies reporting that genetic ablation of *Elob* had minimal effect on mouse embryonic stem cell growth but promoted the growth and survival of a subset of human cancer cell lines (*19*, *27*). We next assessed the consequences of knocking out additional components of the Pol2 SIII elongation complex, ELOA and ELOC, as confirmed by western blotting (**fig. S1G**). Knockout of either *ELOA* or *ELOC* similarly led to a significant reduction in cell number across multiple DMG cell lines (**Fig. 1, J** to **L**), further implicating the Pol2 SIII complex in DMG growth. We also performed BrdU pulse-labeling and cell-cycle analysis in SU-DIPGXIII cells following knockout of *ELOA*, *ELOB*, and *ELOC* and found that *ELOB* loss led to a significant reduction in BrdU(+) S-phase cells and a corresponding increase in the sub-G1 population (**Fig. 1M**). Collectively, these findings suggest a role for ELOB and related Pol2 SIII elongation complex proteins in promoting H3K27M-mutant DMG cell proliferation *in vitro*.

### ELOB promotes H3K27M-mutant DMG tumor growth

To assess whether ELOB is required for DMG tumorigenesis *in vivo*, we initially attempted to generate clonal *ELOB* knockout cell lines; however, we were unable to obtain sufficient cells to support xenograft studies (data not shown). We therefore established orthotopic xenografts of polyclonal control and *ELOB*KO SU-DIPGXIIIP* expressing a ZsGreen/luciferase reporter construct enabling tumor tracking by luciferase imaging. DMG cells were transplanted intracranially into the pons 14 days after sgRNA transduction and selection, and tumor growth was monitored by bioluminescent imaging (BLI). Mice bearing *ELOB*KO tumors generated using two independent sgRNA sequences exhibited reduced tumor luciferase signal compared to controls by day 15 post-implantation (**Fig. 2, A** to **C**), indicating impaired tumor growth upon *ELOB* loss. Given that these studies tracked the growth of polyclonal DMG cell lines composed of a mixture of wildtype and *ELOB*KO cells (**fig. S1F**), we then asked whether unedited ELOB (+) cells might have a growth advantage *in vivo*. We stained brain sections from mice bearing control and *ELOB*KO xenografts for endogenous ELOB and quantified the proportion of ELOB (+) cells. These analyses reveal that most of the control SU-DIPGXIIIP* tumor cells stained positively for ELOB, whereas the ratio of ELOB (+) to ELOB (-) cells increased over time in the *ELOB*KO cohort (**Fig. 2, D** to **F**), suggesting a growth advantage for ELOB (+) cells *in vivo.* Consistently, staining control and *ELOB*KO tumors for MKI67 as a marker of cell proliferation revealed a significant reduction in the proportion of proliferating cells in the *ELOB*KO tumors compared to control xenografts (**Fig. 2, G** and **H**). We also monitored these SU-DIPGXIIIP* xenograft mice for signs of disease progression, including decreased body condition score, hydrocephalus, and other neurological symptoms associated with brain tumor growth, and performed a Kaplan-Meier survival analysis revealing extended survival of mice bearing *ELOB*KO xenografts compared to control tumors (\**p*<0.0295, log-rank test, **Fig. 2I**).

**Figure 2:**
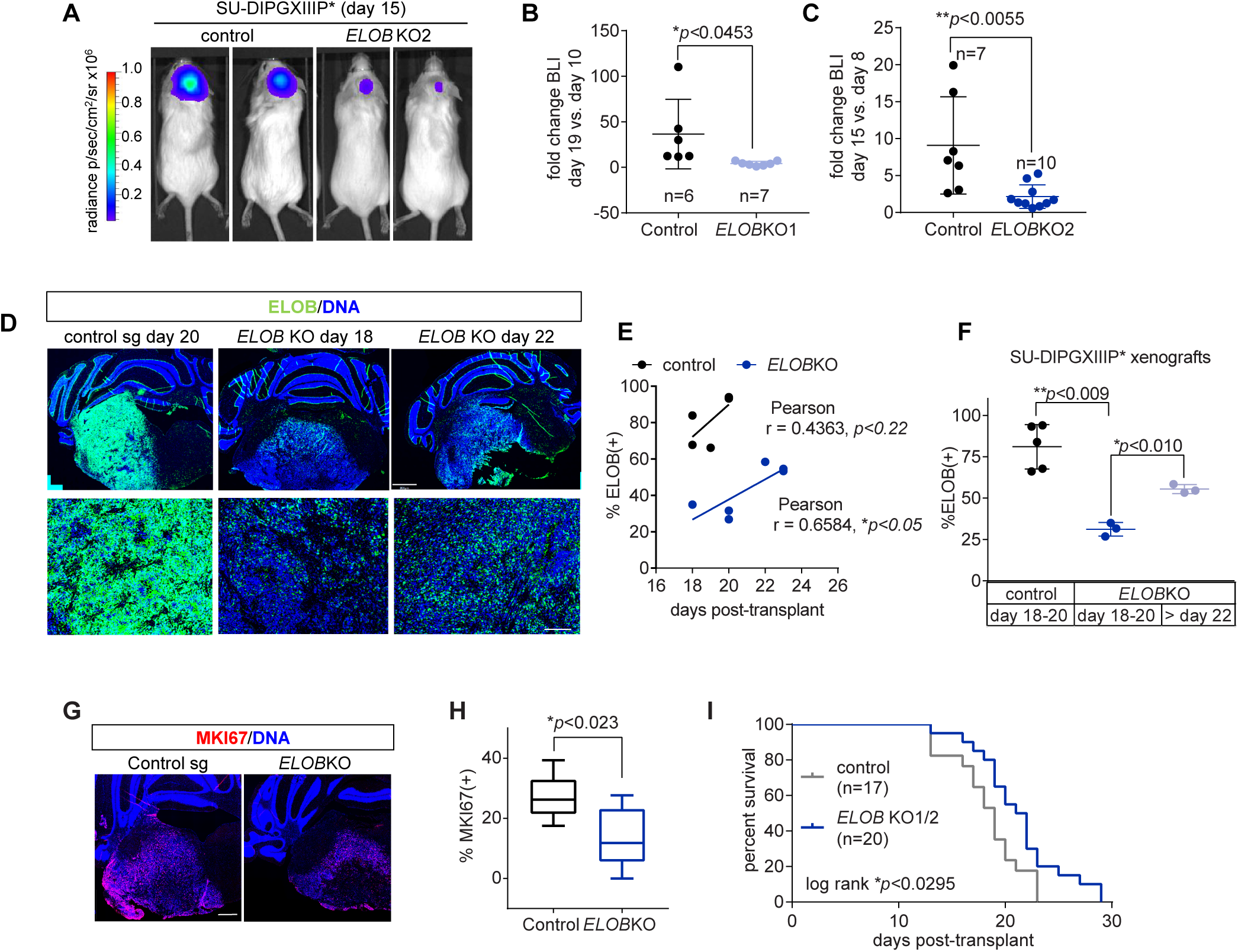
ELOB promotes H3K27M-mutant DMG tumor growth. **(A** to **C)** Representative images **(A)** and quantification **(B** and **C)** from tumor bioluminescent imaging of NSG immunocompromised mice bearing pontine xenografts of control and *ELOB*KO SU-DIPGXIIIP*+Zsgreen/luciferase cells. In panels **(B** and **C)**, two different sgRNA sequences were used to target *ELOB* (KO1 and KO2), and statistical significance was determined using an unpaired Student’s t-test. **(D** to **F)** Representative fluorescence microscopy images of ELOB-stained SU-DIPGXIIIP* xenograft brain sections **(D)** and quantification demonstrating increasing proportions of ELOB(+) cells over time **(E)** and a greater percentage of ELOB(+) cells collected at days 22-23 versus days 18-20 **(F)**. In panel **(D)**, the top scale bar is 800 microns, and the bottom scale bar is 100 microns. **(F** and **G)** Representative images **(F)** and quantification **(G)** of the %MKI67 (+) proliferating cells in control and *ELOB*KO SU-DIPGXIIIP* xenograft tumors. **(H)** Kaplan-Meier survival curve comparing the survival of mice bearing control versus *ELOB*KO SU-DIPGXIIIP* xenograft tumors (*p<0.0295, log-rank test).

### *ELOB*KO causes global disruptions in histone post-translational modifications

We next characterized the consequences of ELOB loss on chromatin and transcriptional regulation in H3K27M-mutant cells by first determining whether *ELOB*KO affects the global abundance of various histone post-translational modifications (PTMs). We isolated histones from control and *ELOB*KO HSJD-DIPG007 and SU-DIPGXIIIP* cells by acid extraction and analyzed them by tandem mass spectrometry, revealing a decrease in the overall abundance of gene-activating histone acetyl marks (H3K18ac, H3K9ac, and H2AXK5ac) and in phosphorylated H3 peptides associated with mitosis (H3S10ph and H3S28ph) in *ELOB*KO versus control cells (**fig. S2A**). These results suggest that ELOB functions to promote or maintain histone PTMs that are associated with active Pol2 transcription and mitotic progression.

Previous studies have reported residual H3K27me2/3 in DMG despite H3K27M-dependent inhibition of Polycomb Repressive Complex 2 (PRC2) activity (*3*, *28*). Given prior studies demonstrating that ELOB and ELOC associate with PRC2 complexes through an interaction with EPOP (*19*, *22*), we next asked if ELOB might regulate global H3K27 modification states. We observe no consistent change in the overall abundance of H3K27 mono-, di-, or tri-methylation or in H3K27ac due to *ELOB* CRISPR-targeting (**fig. S2, B** to **E**). To determine the functional significance of ELOB-PRC2 complexes in promoting DMG growth, we generated *EPOP*KO DMG cell lines to inhibit ELOB recruitment to PRC2 complexes (**fig. S2F**). However, *EPOP* loss of function did not significantly affect the growth and survival of either H3K27M-mutant DMG cells or NHA-hTERT control cells (**fig. S2, G** to **I**). These findings support a role for ELOB in promoting DMG growth independently of its interaction with EPOP and the PRC2 complex, potentially through alternative gene-regulatory complexes like the Pol2 SIII complex.

### ELOB is enriched at genomic loci bound by H3K27M and active histone marks

To further define roles for ELOB in DMG chromatin regulation, we then conducted a series of genome-wide chromatin-profiling studies. We performed CUT&RUN (*29*) using an ELOB-targeting antibody to identify ELOB binding sites in polyclonal control and *ELOB*KO SU-DIPGXIII cells. We analyzed significantly down-regulated peaks in the *ELOB*KO compared to control CUT&RUN data, revealing that ELOB is consistently bound at 3,076 sites (**table S1**). To determine the relationship between ELOB chromatin binding and the incorporation of the H3K27M oncohistone, we performed additional CUT&RUN experiments to identify H3K27M binding sites, revealing 36,229 H3K27M peaks in control DIPGXIII cells where a single copy of H3.1 was targeted (*HIST1H3B* knockout) compared to isogenic H3K27MKO cells lacking the mutant histone generated, as previously described (*30*) (**table S1**). Comparing these ELOB and H3K27M chromatin profiling datasets reveals a co-enrichment of H3K27M at the ELOB binding sites (**Fig. 3, A** and **B**). Additional analyses reveal that 18.3% (6,644/36,229) of the H3K27M binding sites were found within +/- 30 kb of an ELOB binding site, whereas 82% (2,527/3,076) of the ELOB peaks were located within +/- 30 kb of at least one H3K27M peak with a median distance of 0.948 kb (**fig. S3A**). ELOB ChIP-seq in control versus *ELOB*KO SU-DIPGXIII cells further validates a similar overlap between H3K27M CUT&RUN and ELOB ChIP-seq signals and confirms co-occupancy by ELOB and H3K27M at a subset of loci in DMG (**fig. S3B**).

**Figure 3:**
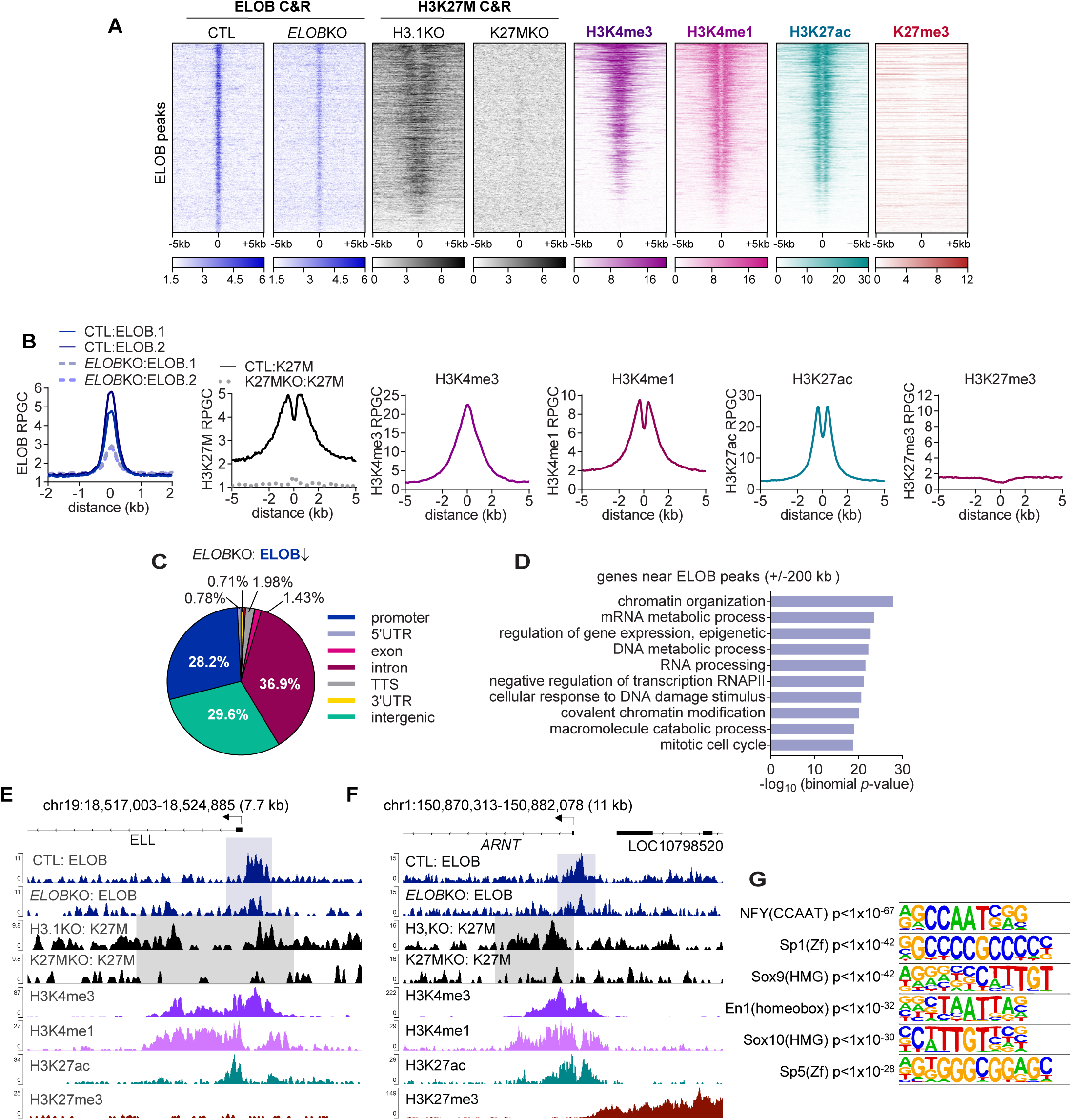
ELOB is enriched at genomic loci bound by H3K27M and enriched in active histone marks. **(A** and **B)** Heatmap **(A)** and average profile plots **(B)** of ELOB binding sites (3,076 peaks) identified by CUT&RUN (C&R), demonstrating co-enrichment in H3K4me3, H3K4me1, H3K27ac, and H3K27M oncohistones and ELOB-bound regions. H3K27M CUT&RUN data was also generated in additional control SU-DIPGXIII cells where a single copy of H3.1 was knocked out by CRISPR-targeting (*HIST1H3B, H3.1KO*) to confirm H3K27M antibody specificity. **(C)** Pie chart showing overlap between ELOB binding sites and various genomic elements, revealing frequent ELOB binding at promoters, intergenic regions, and introns. **(D)** Overrepresented gene categories from Gene Ontology (GO) analysis of the genes near ELOB binding sites (+/- 200 kb) revealing ELOB binding at genes associated with transcription and chromatin regulation, mitosis, and DNA damage pathways. **(E** and **F)** Representative genome browser snapshots showing co-localization of H3K27M (grey boxes), H3K4me1, H3K4me3, and H3K27ac at ELOB peaks (blue boxes) located near the *ELL* **(E)** and *ARNT* **(F)** gene loci. **(G)** Overrepresented transcription factor motifs in the ELOB peak set, including factors involved in glial and brain development like SOX9, SOX10, and EN1 from HOMER motif enrichment analysis.

We then aligned previously generated H3K4me1, H3K27ac, and H3K27me3 ChIP-seq data (*5*) and H3K4me3 CUT&RUN data with these ELOB peaks to assess the baseline chromatin state at these ELOB-bound regions. These integrative analyses reveal an association between ELOB and histone PTMs commonly found at both promoters (H3K4me3) and enhancers (H3K27ac/H3K4me1) (*31*, *32*), but a lack of H3K27me3 enrichment (**Fig 3, A** and **B**). Mapping these ELOB peaks to various genomic elements reveals that ELOB binds at TSS/promoters, gene bodies, and at putative distal enhancers (**Fig. 3C**), consistent with the observed histone PTMs at ELOB binding sites (**Fig. 3, A** and **B**). GO analysis of the genes found near the ELOB peaks (+/-200 kb) indicates that ELOB binds in the vicinity of genes associated with Pol2 transcription, DNA damage response, and mitosis (**Fig. 3D**). As examples, we observe an ELOB peak that overlaps with H3K27M, H3K27ac, and H3K4me1/3 at the *ELL* locus, which encodes a member of the Pol2 super elongation complex (*12*) (**Fig. 3E**), and at the aryl hydrocarbon nuclear transporter (*ARNT*) gene, which has been linked to GBM chemoresistance (**Fig. 3F**) (*33*). Finally, motif enrichment analysis using HOMER reveals an overlap between ELOB chromatin-binding and consensus binding sites targeted by transcription factors involved in glioma cell fate and brain patterning like SOX9, SOX10, and Engrailed (EN1) (**Fig. 3G**) (*34*, *35*). Together, these findings indicate that ELOB co-localizes with H3K27M and active chromatin marks near genes linked to transcriptional regulation and mitosis.

### *ELOB*KO disrupts H3K27me3 and H3K27M chromatin incorporation in DMG

Given the observed co-localization between ELOB and the H3K27M CUT&RUN signal, we then asked whether *ELOB*KO affects H3K27M binding patterns or vice versa through reciprocal loss of function studies. While H3K27MKO had a minimal effect on the ELOB CUT&RUN signal at the top 4,000 ranked H3K27M binding sites characterized by robust oncohistone enrichment (**fig. S3, C** and **D**), *ELOB*KO resulted in a marked reduction in H3K27M binding at 3,402 regions (**table S1**, **Fig. 4, A** and **B**), representing 9.4% (3,402/36,229) of the total H3K27M binding sites identified in these profiling studies. These ELOB-regulated H3K27M peaks co-localize with gene-activating histone marks, like H3K4me3, H3Kme1, and H3K27ac, but lack strong H3K27me3 binding (**Fig. 4, A** and **B**). Following up on previous studies demonstrating that H3K27M-mutant cells exhibit a global reduction in H3K27me3 while preserving sharp peaks of H3K27me3 with reduced heterochromatin spreading at a subset of loci, such as at tumor suppressor genes (*3*, *28*, *36*), we then asked whether the pattern of H3K27me3 chromatin modification might be altered in *ELOB*KO cells. Compared to matched controls, both H3K27MKO and *ELOBKO* resulted in a similar decrease in H3K27me3 signal at 857 regions (**Table S1**, **Fig. 4, C** and **D**). We then assessed the genomic features associated with the ELOB-regulated H3K27M and H3K27me3 peaks, which reveals an overlap with both introns and distal intergenic regions potentially acting as enhancer elements (**Fig. 4E**). These data suggest that ELOB maintains H3K27M and H3K27me3 chromatin binding patterns at a subset of genomic loci in SU-DIPGXIII cells, providing additional evidence for the functional significance of ELOB in sustaining epigenetic dysregulation in DMG.

**Figure 4:**
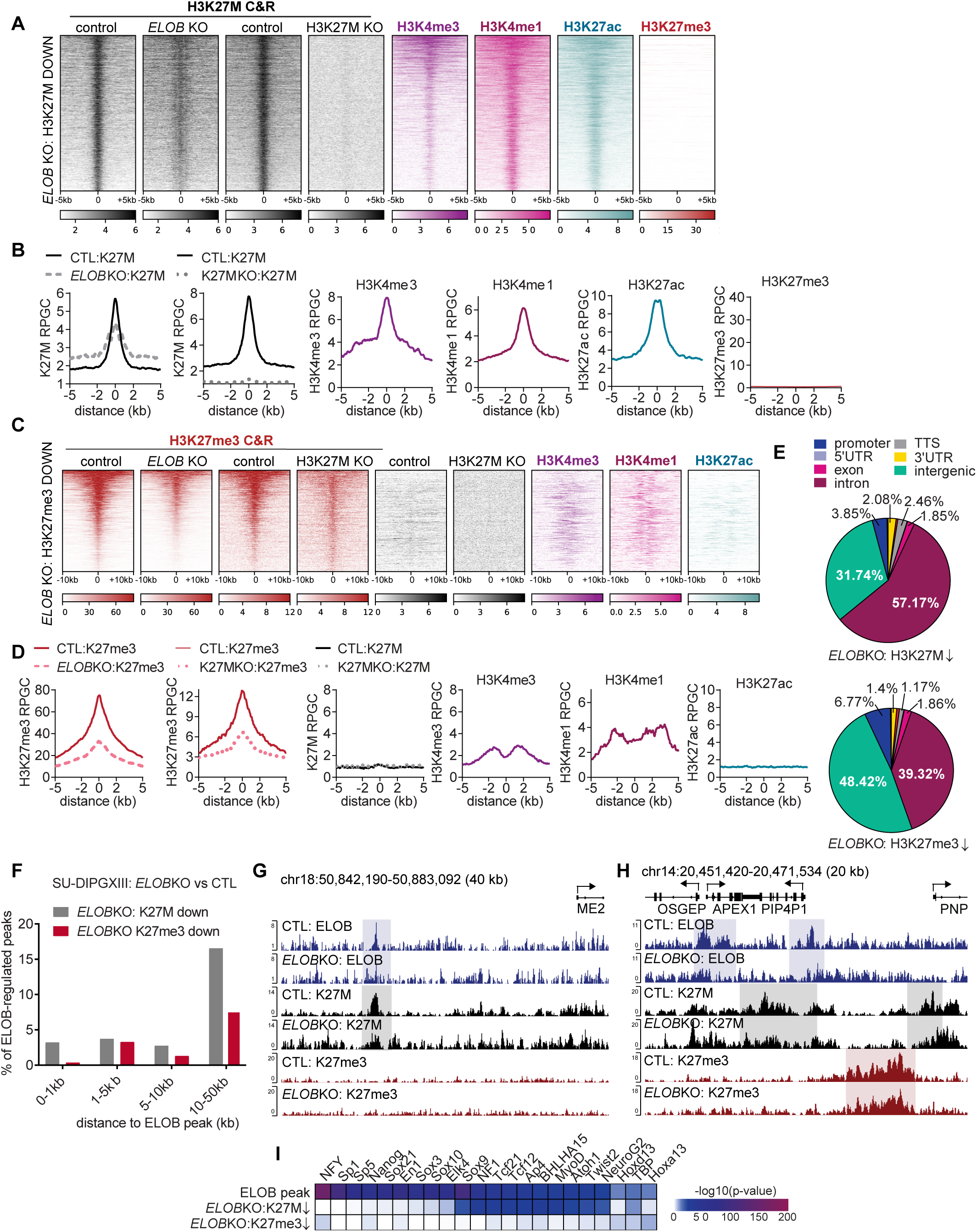
***ELOB*KO disrupts H3K27me3 and H3K27M chromatin incorporation patterns in DMG. (A** and **B)** Heatmaps **(A)** and average profile plots **(B)** centered on genomic regions where *ELOB*KO reduced H3K27M binding (3,042 sites). H3K4me3 and H3K27me3 CUT&RUN data and H3K27ac and H3K4me1 ChIP-seq data are also shown, demonstrating co-localization between these H3K27M-bound sites with H3K4me1, H3K4me3, and H3K27ac, but not H3K27me3. **(C** and **D)** Heatmaps **(C)** and average profile plots **(D)** centered on regions where both H3K27M loss and *ELOB*KO reduced H2K27me3 CUT&RUN signal (857 sites). Alignment with other ChIP-seq and CUT&RUN datasets, as in panels **(A** and **B)**, demonstrates co-localization of these H3K27me3-down peaks with the enhancer marker, H3K4me1, and the promoter marker, H3K4me3, but low H3K27ac enrichment. **(E)** Overlap between differential H3K27M peaks (top graph, see panels **A** and **B**) and differential H3K27me3 peaks (bottom graph, see panels **C** and **D**) and various genomic elements suggesting localization to promoters, introns, and intergenic regions. **(F)** Histogram showing the distance between the *ELOB*KO-downregulated H3K27M peaks (grey bars) and –downregulated H3K27me3 peaks (red bars), revealing reduced distances between differential H3K27M peaks and ELOB binding sites. **(G** and **H)** Genome browser snapshots showing ELOB binding (blue boxes) upstream of the *malic enzyme 2* (*ME2)* gene locus corresponding with an ELOB-regulated H3K27M peak (grey box) **(G)** and ELOB binding corresponding with regional changes in H3K27M (grey box) and H3K27me3 (red box) near the *OSGEP/APEX1/PIP4P1* gene loci **(H)**. **(I)** Heatmap of –log_10_(*p*-values) for enriched transcription factor binding motifs at ELOB binding sites and at ELOB-regulated H3K27M and H3K27me3 peaks, as identified by HOMER.

We then asked whether *ELOB*KO might have either direct or indirect effects on local H3K27M or H3K27me3 binding patterns by assessing whether ELOB peaks are localized in the same gene loci as the *ELOB*KO-altered H3K27M and H3K27me3 peaks. We determined the distance between the ELOB-regulated H3K27M peaks and H3K27me3 peaks, revealing that 26% of the *ELOB*KO down-regulated peaks were within 50 kb of an ELOB binding site (889/3,402 peaks), whereas 12% of the reduced H3K27me3 peaks were within 50 kb of an ELOB binding site (103/857) (**Fig. 4F**). These findings indicate that ELOB loss results in both local and indirect effects on H3K27M and H3K27me3 chromatin incorporation patterns. For example, we observed co-localization between H3K27M and ELOB at a region upstream of the *Malic Enzyme 2* (*ME2*) transcription start site (TSS) and found that H3K27M chromatin binding was reduced at this site following *ELOB*KO (**Fig. 4G**). We also observe localized changes in ELOB, H3K27M, and H3K27me3 chromatin enrichment in the vicinity of the *APEX1, PNP, SUPT16H,* and *RPGRIP1* gene loci, but the ELOB-regulated H3K27M and H3K27me3 peaks did not directly overlap with the ELOB binding sites (**Fig. 4H** and **fig****. S3E**). These results suggest that ELOB may contribute to locus-specific modulation of H3K27M and H3K27me3, both at directly co-occupied sites and at nearby regions.

Further GO analysis on the genes nearest the *ELOB*KO-altered H3K27M and H3K27me3 peaks reveals that *ELOB*KO regulates H3K27M and H3K27me3 binding at genes associated with Pol2 transcription, cell proliferation, and central nervous system development (**fig. S3, F** and **G**), further highlighting ELOB-dependent control of chromatin architecture at genes associated with key biological processes underlying gliomagenesis. Transcription factor motif calling reveals an enrichment in binding sites for SOX9, MYOD1, ATOH1, and NEUROG2 among both the consensus ELOB binding sites and the ELOB-regulated H327M peaks (**Fig. 4I**). In contrast, the ELOB-regulated H3K27me3 peaks were weakly associated with unique transcription factor motifs, including HOXA13 and HOXD13 binding sites (**Fig. 4I**). Overall, these results support a model whereby ELOB cooperates with H3K27M to promote aberrant chromatin states via altered H3K27M and H3K27me3 incorporation at a subset of gene loci linked to DMG malignancy.

### *ELOB*KO alters Pol2 transcriptional activity at bivalent chromatin regions to control developmental and metabolic genes

To define ELOB’s role in DMG gene regulatory programs, we performed both poly (A) RNA-seq and PRO-seq (Precision Run-On sequencing) on control and ELOBKO cells, enabling both high-resolution mapping of the location and activity of transcriptionally-engaged Pol2 across the genome and analysis of changes in mRNA transcript abundance. RNA-seq analysis on SU-DIPGXIII and BT245 DMG cells identified 1,625 up- and 648 down-regulated genes following *ELOB*KO (*padj<0.05, ≥1.4 fold change, **table S2**, **Fig. 5A**, **fig. S4, A** and **B**). Complementary PRO-seq analysis reveals widespread alterations in nascent transcription with decreased Pol2 transcription at 1,084 genes and increased Pol2 activity at 1,183 genes (**table S3**). Consistent with these findings, *ELOB*KO resulted in increased PRO-seq signal in both the promoters and gene bodies of the upregulated genes (**Fig. 5B**), demonstrating an expected association between Pol2 transcriptional activity and transcript abundance. These transcriptional changes are consistent with our analysis of chromatin alterations induced by *ELOB*KO, which include remodeling of both H3K27-associated active chromatin regions and the repressive H3K27me3 mark (**Fig. 4** and **S3**).

**Figure 5.**
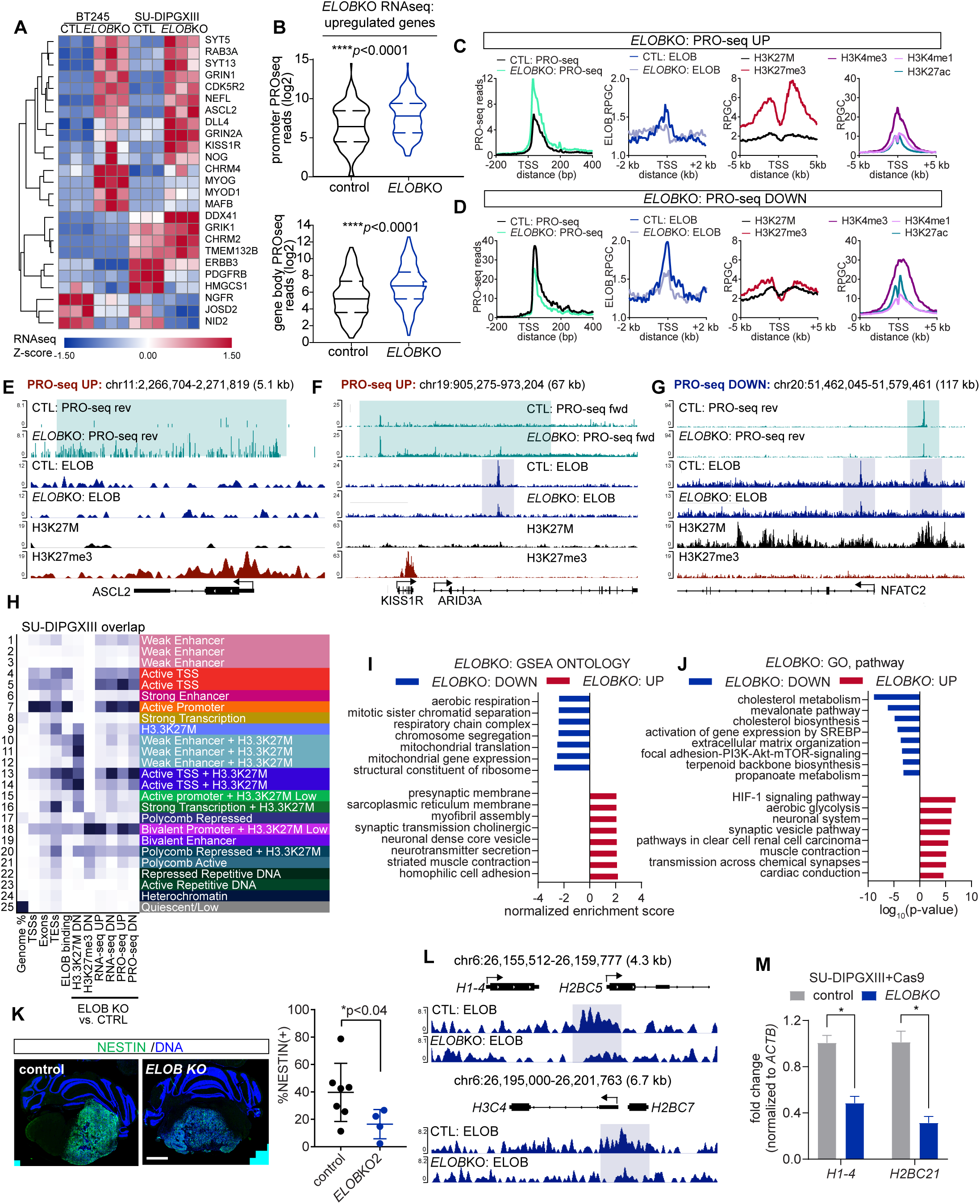
***ELOB*KO alters Pol2 transcriptional activity at bivalent chromatin regions to control developmental and metabolic genes.** (**A**) Heatmap showing Z-scores from RNA-seq analysis of control and *ELOB*KO SU-DIPGXIII and BT245 cells. See also **table S4. (B)** Quantification of PRO-seq reads at *ELOB*KO-upregulated genes in SU-DIPGXIII cells demonstrating a corresponding increase in Pol2 activity at both the gene promoters (top panel) and gene bodies (bottom panel) due to *ELOB* loss. **(C** and **D)** Average PRO-seq profiles and CUT&RUN signal centered on transcription start sites (TSS) with increased **(C)** or decreased **(D)** PRO-seq reads in *ELOB*KO cells. See also **table S3**. **(E** to **G)** Genome browse snapshots revealing decreased PRO-seq signal at the H3K27me3-enriched *ASCL2*, *KISS1R,* and *ARID3A* gene loci with regional ELOB and H3K27me3 enrichment **(E** and **F)** and decreased PRO-seq signal at the *NFATC2* gene, which is bound by both ELOB and H3K27M **(G)**. Differential PRO-seq peaks and ELOB-binding regions are marked by green and blue boxes. **(H)** Heatmap of ChromHMM results integrating ChIP-seq, CUT&RUN, RNA-seq, and PRO-seq data revealing dynamic Pol2 activity (PRO-seq-UP and -DN), enhancer signatures, and bivalent H3K27me3/H3K4me3 chromatin at ELOB-regulated targets. **(I** and **J)** GSEA **(I)** and GO analysis **(J)** of *ELOB*KO RNA-seq data showing downregulation of genes associated with cell division, mitochondria, and metabolism, and upregulation of genes associated with neuronal function and excitatory tissues. GSEA results represent normalized enrichment scores (NES) for non-redundant gene sets (*FDR<*0.01). **(K)** Representative images of NESTIN staining (left panel) and quantification of the %NESTIN(+) (right panel) stem/progenitor-like cells in control and *ELOB*KO SU-DIPGXIIIP* xenografts. Scale bar is 800 microns. **(L)** Genome browser snapshots showing ELOB binding at histone genes. **(M)** RT-PCR quantification of *H1-4* and *H2BC21* transcript expression normalized to *ACTB* from control versus *ELOB*KO SU-DIPGXIII cells.

To determine whether altered Pol2 activity correlates with ELOB binding, we aligned the ELOB CUT&RUN data with the TSS of genes exhibiting either increased or decreased PRO-seq signal but detected minimal ELOB enrichment at these sites (**Fig. 5**, **C** and **D**). Focusing on the genes that were up-regulated by *ELOB*KO, we also calculated the Pol2 pausing index, which was not significantly changed in the *ELOB*KO versus control cells (**fig. S4C**), indicating that ELOB loss does not broadly affect Pol2 occupancy at the TSS or promoter-proximal pausing. These data suggest that ELOB binding at TSS may be transient and difficult to capture by CUT&RUN, or alternatively, that ELOB-dependent modulation of Pol2 dynamics is not limited to the proximal promoter region. Consistent with this model, *ELOB*KO resulted in increased PRO-seq signal at the *ASCL2, KISS1R, ARID3A,* and *GSX1* gene loci (**Fig. 5**, **E** and **F**, and **fig. S4D**) and decreased PRO-seq signal at the *NFATC2, ACSS1* (*Acyl-CoA Synthetase Short Chain Family Member 1*), and *FAM72A* gene loci (**Fig. 5G**, **fig. S4**, **E** and **F**). Examination of ELOB binding at these target genes reveals ELOB enrichment both near TSS/promoters (*NFATC2*, *ACSS1 FAM72A;* **Fig. 5G** and **fig****. S4**, **E** and **F**) as well as in gene bodies (*ARID3A*, *NFATC2;* **Fig. 5, F** and **G**). Collectively, these analyses suggest potentially complex roles for ELOB in regulating Pol2 dynamics at promoter-proximal and genic regions.

To explore a potential association between ELOB-dependent changes in Pol2 transcription and aberrant DMG chromatin states, we examined H3K27M occupancy and histone modification patterns at genes with altered PRO-seq signal in *ELOB*KO versus control cells. We find that *ELOB*KO results in increased Pol2 transcriptional activity at genes marked by high H3K27me3 and moderate H3K27M binding (**Fig. 5C**), whereas genes with decreased Pol2 activity were marked by relatively lower H3K27me3 but increased H3K27M and active histone mark enrichment (**Fig. 5D**). Consistent with the average profile plots, we observe H3K27me3 enrichment at the *ASCL2, KISS1R, and GSX1* gene loci accompanied by an increase in the PRO-seq signal following *ELOB* loss (**Fig. 5, E** and **F**, and **fig. S4D**), whereas H3K27M binding was observed at the *NFATC2*, *ACSS1*, and *FAM72A* genes, which exhibited reduced PRO-seq signal in the ELOBKO cells (**Fig. 5G**, **fig. S4**, **E** and **F**).

To more systematically assess correlations between ELOB and H3K27M-dependent chromatin and transcriptional dysregulation, we applied ChromHMM (*37*) to integrate chromatin profiling, RNA-seq, and PRO-seq data generated here with published datasets to derive a 25-state model of chromatin and ELOB-dependent transcriptional regulation in DMG (**fig. S4G**). This chromatin state model encompasses a variety of gene regulatory states, including gene silencing (H3K27me3-enriched), gene activation (TSS, enhancers), and bivalent chromatin (H3K27me3/H3K4me3). Overlaying the *ELOB*KO multiomic data onto this framework reveals that both ELOB binding sites and ELOB-regulated H3K27M peaks preferentially coincide with promoter and enhancer states (**Fig. 5H**). In addition, ELOB-dependent changes in histone modifications, PRO-seq signal, and gene expression were associated with a bivalent (H3K27me3/H3K4me3) chromatin state, which has been linked to DMG malignancy and the regulation of stemness and differentiation in development (**Fig. 5H**) (*38*, *39*). Taken together, these analyses suggest that ELOB regulates chromatin states and Pol2 dynamics at subsets of genes marked by either H3K27M or H3K27me3 enrichment, which are characteristically dysregulated in H3K27M mutant-DMG, further reinforcing a role for ELOB in chromatin and transcriptional dysregulation in these brain tumors.

GO analysis and GSEA on the differentially expressed genes (DEGs) in the *ELOB*KO cells suggest that *ELOB*KO silences the expression of transcripts associated with critical biological processes, such as metabolism, ribosome biogenesis, and mitosis, whereas transcripts associated with neuronal activity or expressed in other excitatory cells, like muscle and cardiac tissue, were upregulated upon *ELOB* loss (**Fig. 5**, **I** and **J**). These findings point to a role for ELOB in the regulation of genes linked to cell growth and differentiation states in brain tumors. Staining of SU-DIPGXIIIP* xenograft tumors confirms that the neural progenitor cell marker, Nestin (NESTIN), was reduced in *ELOB*KO tumors, suggesting that ELOB regulates DMG cell phenotypes *in vivo* (**Fig. 5K**). We also observe ELOB binding at histone gene loci in both our CUT&RUN analysis and in subsequent ChIP-qPCR validation studies (**Fig. 5L** and **fig****. S4H**) and a corresponding decrease in histone transcript expression by RT-qPCR (**Fig. 5M**), suggesting a role for ELOB in histone gene transcription and biogenesis. Finally, *ELOB*KO alters the expression of TGFβ and WNT signaling genes, including *Wntless* (*WLS*), which is required for WNT secretion (**fig. S4I**). Follow-up studies reveal that a majority of DMG cell lines are sensitive to inhibitors of WNT secretion (LGK974 and WNT-C59; **fig. S4**, **J** and **K**) (*40*, *41*), confirming that optimal DMG growth depends on an ELOB-regulated pathway. Together, these results demonstrate that ELOB coordinates transcriptional programs essential for DMG proliferation and lineage specification, including histone genes and WNT signaling genes.

### ELOB and ELOB-binding proteins are overexpressed in high grade glioma and correlate with poor clinical outcomes

In parallel to our functional studies revealing roles for ELOB in controlling DMG growth, chromatin states, and gene regulation, we analyzed the expression of various Pol2 elongation factors in publicly available, pediatric brain tumor patient transcriptomic and proteomic datasets (*42*, *43*). We first asked whether there were any differences in *elongin* mRNA expression in pediatric low (LGG) and high grade glioma (HGG) samples grouped by histone mutation status. This analysis finds increased *ELOA* expression in HGG samples regardless of histone mutation status compared to LGG, but no significant differences in *ELOC* and *ELOB* across these diagnostic groups (**fig. S5**, **A** to **C**). Further Kaplan-Meier survival analyses comparing patients with high versus low *elongin* mRNA expression reveal a correlation between increased *ELOB* and *ELOC* transcripts and reduced survival time in pediatric HGG (**Fig. 6**, **A** and **B**) but no significant correlation between *ELOA* transcript abundance and patient outcomes (**fig. S5D**). We then characterized ELOB and ELOC protein expression in a publicly available pan-pediatric brain tumor patient proteomics dataset (CPTAC) grouped by brain tumor diagnostic subtype. This analysis suggests increased expression of ELOB in high grade glioma (HGG), atypical teratoid rhabdoid tumors (ATRT), medulloblastoma (MB), and ependymoma (EPM) compared to low grade glioma (LGG) at a protein level (**Fig. 6C**). ELOC was similarly highly expressed in HGG, ATRT, and EPM compared to LGG samples (**Fig. 6D**).

**Figure 6.**
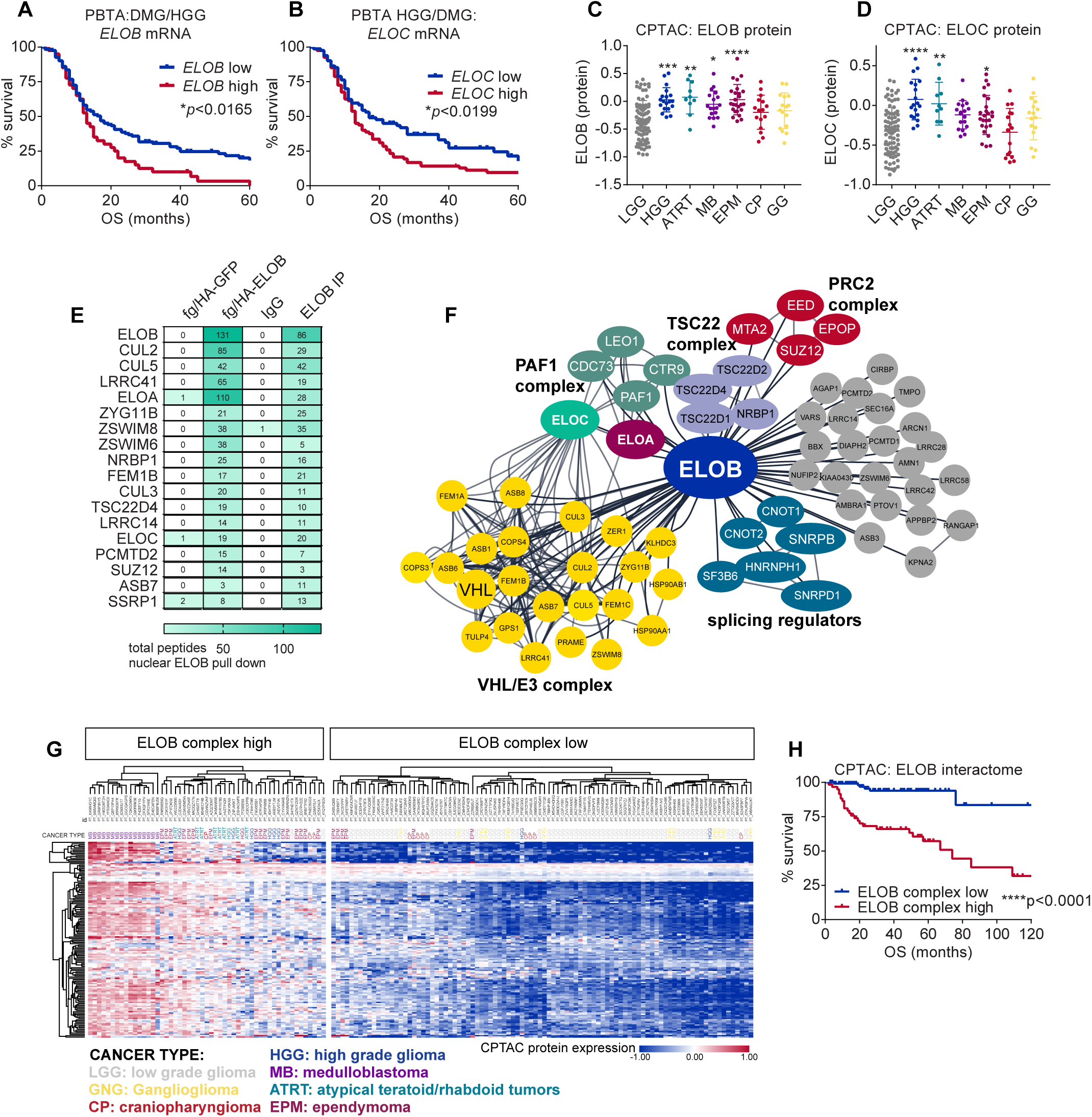
ELOB and ELOB-binding proteins are overexpressed in pediatric high grade glioma and correlate with poor clinical outcomes. **(A** and **B)** Kaplan-Meier curves showing survival of HGG/DMG patients grouped into high and low *ELOB* (A) *and ELOC* (B) transcript expression based on publicly available patient tumor RNA-seq data (PedcBio) with statistical significance determined by the log-rank test (\**p*<0.05). **(C** and **D)** Relative protein expression of ELOB **(C)** and ELOC **(D)** in patient data from a pan-pediatric brain tumor proteomics cohort (CPTAC) grouped by tumor subtype (LGG = low grade glioma, HGG = high grade glioma, ATRT = atypical teratoid rhabdoid tumors, MB = medulloblastoma, EPM = ependymoma, CP = craniopharyngioma, GG = ganglioglioma). Statistical significance was determined by one-way ANOVA comparing all samples to LGG with \**p*<0.05, \*\**p*<0.01, \*\*\**p*<0.001, and \*\*\*\**p*<0.0001. **(E)** Table showing total peptides from LC-MS/MS quantification of nuclear pulldowns of either flag/HA-ELOB from BT869 cells or endogenous ELOB from SU-DIPGXIII cells with fg/HA-GFP and IgG pulldowns serving as negative controls. See also **table S4**. **(F)** ELOB interactome diagram from overlap between flag/HA-ELOB and endogenous ELOB protein complex isolation showing interactions with transcriptional regulators, E3 ubiquitin ligase complexes, splicing regulators, and Polycomb (PRC2) complex members. **(G)** Heatmap showing expression of ELOB protein-protein interactome in pediatric brain tumor patient proteomics data grouped by high and low expression of ELOB-associated proteins. **(H)** Kaplan-Meier survival curves indicating decreased survival time in patients with tumors expressing high levels of ELOB and associated proteins in a pan-pediatric brain tumor analysis (\*\*\*\**p*<0.0001, log-rank test).

To more broadly assess potential correlations between ELOB gene-regulatory activity and pediatric glioma patient outcomes, we derived an ELOB protein-protein interactome combining analysis of proteins associated with either endogenous ELOB or flag-ELOB by immunoprecipitation and mass spectrometry (**table S4**, **Fig. 6**, **E** and **F**). We identified numerous ELOB-interacting partners, including components of VHL/E3 ubiquitin ligase complexes (e.g. ZSWIM6,8 and ZYG11B; shown in yellow), the PAF1 complex (shown in green), and the Polycomb (PRC2) complex (shown in red) (**Fig. 6F**). We then clustered the pan-pediatric glioma patient proteomics data into two groups exhibiting either high or low expression of ELOB-interacting proteins (**Fig. 6G**), which suggests that ELOB-associated proteins are highly expressed in HGG samples and other brain tumors like medulloblastoma and ependymoma compared to LGG samples (shown in grey). Consistently, many ELOB-interacting proteins are upregulated at the mRNA level in H3 wildtype, H3K27M-mutant, and H3G34R/V-mutant patient HGG samples compared to LGG (**fig. S5**, **E** to **L**). Finally, a Kaplan-Meier survival analysis comparing the ELOB complex-high and -low tumors reveals a correlation between increased ELOB interactome expression and reduced pediatric brain tumor patient survival (**Fig. 6H**). Overall, these findings suggest an association between altered ELOB protein complex expression and a broader subset of malignant brain tumors not limited to gliomas bearing the H3K27M mutation, providing further evidence that Elongin-associated Pol2 and chromatin regulatory pathways may contribute to brain tumor development and progression.

## DISCUSSION

Dysregulation of transcription is a hallmark of H3K27M-mutant DMG and other brain tumors and a potential therapeutic vulnerability (*7*, *11*, *44*, *45*). Several Pol2 co-factors have been identified as drivers of glioma growth and survival (*7–11*), yet our knowledge of the mechanisms and molecular effectors maintaining aberrant gene networks in glioma and their context-dependent roles in H3K27M-mutant DMG development and progression remains incomplete. We have identified multiple novel genes associated with Pol2 elongation regulation as genetic dependencies in DMG, including ELOB and other members of the Pol2 SIII complex (*15*). Our further studies reveal that ELOB promotes DMG growth *in vitro* and *in vivo*, implicating ELOB and the Pol2 SIII complex in DMG disease biology and malignancy.

Our chromatin profiling studies reveal an overlap between ELOB binding sites and H3K27M oncohistone-enriched regions, suggesting a potential role for ELOB in sustaining aberrant DMG gene regulatory programs rather than broad functionality in maintaining Pol2 transcriptional activity on a genome-wide scale. These findings are consistent with previous studies reporting that ablation of specific Pol2 elongation regulators affects the expression of select transcripts, such as genes involved in cell fate specification and differentiation (*19*, *23*, *46–49*) and genes involved in cellular response to heat shock and other stressors (*47*, *50–52*). Consistently, we find that ELOB binding sites are enriched in motifs targeted by transcription factors involved in glial cell development, like SOX9 and SOX10, and our further RNA-seq and PRO-seq studies link ELOB to the transcription of genes expressed in excitatory tissues like neurons and muscle cells as well as metabolic and proliferation genes sustaining cancer growth. Increased Pol2 recruitment to histone gene loci occurs across multiple solid tumor types and was recently shown to correlate with poor clinical outcomes in cancer (*53*). Our further analyses reveal that ELOB binds histone gene loci and regulates histone gene expression. These results establish ELOB as a regulator of histone biogenesis, which is likely under high demand in proliferative cancer cells like DMG. Finally, we find that metabolic genes involved in glycolysis, mitochondrial function, and cholesterol metabolism were dysregulated by *ELOB*KO, consistent with our recent work and other studies linking transcriptional regulators like the SAGA complex to the control of metabolic gene expression in DMG (*24*). Overall, these results suggest that ELOB orchestrates the chromatin state and expression of key transcriptional, metabolic, and proliferation genes that drive DMG growth and tumorigenesis.

Our chromatin profiling studies reveal that *ELOB*KO results in a loss of H3K27M and H3K27me3 chromatin binding at a subset of loci, whereas ELOB binding is largely unchanged in isogenic cell lines in which the H3K27M-mutant allele was removed by CRISPR targeting. Additional analyses suggest a role for ELOB in promoting Pol2 activity at H3K27me3-silenced bivalent genes and in repressing Pol2 transcription in H3K27M-associated genes, consistent with a partial reversal of DMG-associated chromatin and transcriptional states upon ELOB loss. These findings challenge the prevailing assumption that chromatin state predominantly drives the recruitment of transcriptional regulators by identifying a Pol2 regulator that regulates H3K27M occupancy at a subset of cis-regulatory elements.

A limitation of our current study is that the mechanisms by which *ELOB*KO leads to altered H3K27M and H3K27me3 incorporation at a subset of loci are not yet known. Recent structural studies highlight roles for ELOB-associated Pol2 elongation regulators, including Spt6, the PAF1 complex, and the SIII complex, in regulating nucleosome retention and transfer as Pol2 transcribes through chromatin, suggesting that ELOB loss may alter H3K27M binding patterns through disruptions in transcription-coupled nucleosome recycling and assembly (*25*, *54–56*). Alternatively, *ELOB*KO may regulate H3K27M incorporation via indirect mechanisms involving altered H3.3/H3.1 replacement across cell divisions (*57–59*). A third possibility is that ELOB may modulate H3K27M binding via enhancer-promoter interactions, given that ELOB’s binding partner, ELOA, has been linked to enhancer regulation (*20*). Future studies may shed light on how ELOB and other Pol2 elongation regulators control H3K27M chromatin binding patterns and associated epigenetic defects on a deeper mechanistic level.

Finally, our correlative analysis of publicly available pediatric brain tumor proteomic and transcriptomic data reveals increased expression of a subset of ELOB-binding proteins, not only in H3K27M-mutant DMG but also in other pediatric cancer subtypes like medulloblastoma and ATRT. These results suggest that ELOB is not a specific dependency in tumors driven by H3K27M oncohistones but may instead be co-opted to maintain aberrant Pol2 in a subset of cancers with diverse genetic and epigenetic drivers. Recent studies have reported roles for ELOB in breast cancer, melanoma, prostate cancer, and other malignancies, and for ELOA in gastric cancer and colorectal cancer (*27*, *60*, *61*). Additional studies may delineate the specific functions of Pol2 elongation regulators across pediatric and adult cancers, which may uncover additional cancer dependencies on specific Pol2 regulatory complexes and help prioritize tumor contexts most likely to benefit from Pol2-targeting therapies.

## MATERIALS AND METHODS

### Cell lines and culture conditions

Patient-derived H3K27M DMG cell lines (SU-DIPGVI, SU-DIPGXIII, SU-DIPGXIIIP*, BT245, and BT869 cells) were kindly provided by Dr. Michele Monje and Dr. Keith Ligon and cultured in suspension as gliomaspheres in serum-free media as previously described (*4*). Cell lines expressing Cas9, pHIV Zsgreen/luciferase, and various sgRNAs cloned into pLenti_SpsmB1_sgRNA_Hygro were generated by lentiviral transduction as previously described (*5*).

### CRISPR drop-out screen

SU-DIPGVI and SU-DIPGXIII cells expressing Cas9 were infected with a chromatin-focused sgRNA lentiviral library at a coverage of 1,500 cells per sgRNA in triplicate. Two days after infection, cells were selected for 3 days with 2 μg/mL puromycin (Gold Biotechnology, P-600) and then allowed to grow for an additional three days. Reference samples were collected to quantify the initial sgRNA abundance. The remaining infected cells were outgrown for 5 weeks, and surviving cells were harvested. Genomic DNA was isolated from all samples, and the sgRNA sequences were amplified by PCR before sequencing on a HiSeq2500 sequencer (Illumina). STARS was used to score depleted genes under each condition relative to the reference samples (https://portals.broadinstitute.org/gpp/public/software/stars).

### Western blotting and antibodies

Whole cell lysates were prepared using RIPA lysis buffer (50 mM Tris-HCl pH 8.0, 0.1% SDS, 1% NP-40, 0.5% Na deoxycholate, and 150 mM NaCl) with protease (Millipore Sigma, 5892970001) and phosphatase inhibitors (Millipore Sigma, 4906837001) in 1xLaemmli buffer containing 2.5% v/v β-mercaptoethanol. Samples were resolved on SDS-PAGE gels, transferred to nitrocellulose using Towbin’s buffer, and then blocked with 4% milk dissolved in PBS with 0.1% Tween-20. Westerns were performed using the following primary antibodies: ELOB (Abcam, ab45297), total H3 (Abcam, ab1791), EPOP (ActiveMotif, C01189), ELOA (Cell Signaling Technology, 3685), ELOC (Bethyl, A300-943A), or ACTB (Millipore Sigma, A5441), followed by incubation with HRP-conjugated secondary antibodies (Sino Biologica, SSA004-200) and ECL detection.

### Cell proliferation and viability assays

Cell proliferation assays were performed by seeding 1×10^5^-5×10^5^ cells in 12- or 6-well plates at various time points after selection for sgRNA-expressing cells. Cells were counterstained with 0.02% Trypan Blue, and live, unstained cells were counted using a hemocytometer. For CellTiter-Glo assays, 10,000 DMG cells/well were seeded in 96-well plates and exposed to serial dilutions of LGK974 (MedChemExpress, HY-17545) or WNT-C59 (MedChemExpress, HY-15659) and quantified using the CellTiter-Glo Luminescent Cell Viability Assay (Promega, G7573) after 7 days of treatment. Luminescence was measured on a PerkinElmer VICTOR X3 multilabel plate reader and normalized to DMSO-treated controls.

### Immunofluorescence staining

Cas9 (+) DMG cells transduced with control or *ELOB* sgRNAs were plated on laminin-coated coverslips and fixed with 4% PFA for 20 minutes prior to staining. After washing in PBST (PBS with 0.1% Tween20), cells were permeabilized with 0.25% TritonX-100 in PBS for 30 minutes and blocked in PBST containing 10% FBS. Coverslips were then incubated in block (5% FBS in PBST) containing 1:5000 anti-ELOB (Abcam, ab45297). After washing 4x with PBST, the coverslips were incubated with Alexa Fluor-conjugated secondary antibodies (ThermoFisher Scientific, A11008 and A11012) for 30 minutes at room temperature and then washed 4x with PBST before counterstaining DNA with Hoechst and mounting with Prolong Gold (ThermoFisher Scientific, P36390) on glass slides.

### Xenograft models of DMG tumor growth

NOD SCID gamma (NSG) mice (Jackson Laboratory, 005557) were used for all *in vivo* experiments. Orthotopic xenografts were generated by implanting 5×10^5^ SU-DIPGXIIIP* or BT245 cells engineered to express Cas9, ZsGreen/luciferase, and either control or ELOB-targeting sgRNAs into the pons of 6–8-week-old mice at the following Lambda coordinates: x=-1 mm, y=- 0.8 mm, z=-5 mm. Tumor progression was assessed using the IVIS Spectrum In Vivo Imaging System (PerkinElmer). For bioluminescence imaging, mice received a subcutaneous injection of D-luciferin potassium salt (75 mg/kg in sterile PBS; Promega, E1605) and were anesthetized with 2% isoflurane in medical air. Serial images were collected using the automated exposure settings. Peak bioluminescence signal intensity within defined regions of interest (ROI) was quantified using Living Image Software and reported as photon flux (p/sec/cm²/sr). Representative planar bioluminescence images are shown with the indicated adjusted minimum and maximum thresholds.

### Brain tumor tissue collection and IHC

Mice were perfused with saline followed by 4% paraformaldehyde (PFA), and brain tissue was dissected and fixed in 4% PFA overnight, followed by storage in 70% ethanol prior to staining to detect ELOB (Abcam, ab45297), NESTIN (Cell Signaling Technology, 33475), or MKI67 (Abcam, ab15580). Representative images were analyzed using QuPath and ImageJ.

### CUT&RUN and ChIP-seq analysis

CUT&RUN and ChIP-seq were conducted as previously described in biological duplicates (*5*, *33*) using the following antibodies: ELOB (Abcam, ab45297), H3K4me3 (Abcam, ab8580), H3K27M (Millipore Sigma, ABE419), H3K27me3 (Millipore Sigma, 07-449), and IgG (Cell Signaling Technology, 2729S). Paired-end libraries for sequencing were prepared using the NEBNext Ultra II DNA Library Prep Kit for Illumina (New England Biotechnology, E7645S) and sequenced on an Illumina HiSeq2500. CUT&RUN sequencing data were aligned to hg38 using bowtie2 after removing duplicates using TrimGalore (*61*). MACS2 (*62*) was used to call peaks compared to IgG control, and significant ELOB and H3K27M and H3K27me3 peaks in the *ELOB*KO and H3K27M KO cells were assessed using DiffBind (*63*). ChIP-seq data from a previous study(*5*) using an H3K27ac antibody (ActiveMotif, 39133) and an H3K4me1 antibody (Abcam, ab8895) to enrich for these chromatin marks were also included in the integrative analyses. Motif analysis was done using HOMER to identify enriched motifs within differential peak sets. ChIP-PCR was conducted as previously described using the following antibodies: ELOB ab1, Abcam, ab45297; ELOB ab2, Bethyl, A304-008A; or IgG: Cell Signaling Technology, 2729.

### PRO-seq and RNA-seq analysis

PRO-seq library construction and data analysis was conducted in triplicate on 1×10^6^ control and *ELOB*KO SU-DIPGXIII permeabilized cells as previously described(*62*). For RNA-seq studies, equal numbers of control and *ELOB*KO SU-DIPGXIII and BT245 cells were collected for analysis. Total RNA was purified using TRIzol (ThermoFisher Scientific, 15596018) and mRNAs were isolated using the NEBNext poly (A) mRNA isolation module (New England Biotechnology, E7490L). mRNAs were fragmented by heating to 95°C for ten minutes in a thermocycler and then libraries were generated using the NEBNext Ultra Directional RNA Library Prep Kit for Illumina (New England Biotechnology, E7420L). Multiplexed libraries were pooled in equimolar ratios and were purified from a 2% TAE-agarose gel prior to sequencing to a length of 50 bases using an Illumina HiSeq2500. RNA-seq reads were selected for quality and length using trimgalore and then aligned to hg38 using STAR. Gene level counts were generated using the Featurecounts package and the resulting counts were analyzed in R using DeSEQ2 to identify differentially expressed genes. Genes that were up- or down-regulated at least 1.4 fold with an padj.<0.05 were considered differentially expressed for downstream analyses.

### ELOB protein complex and histone isolation and analysis by LC-MS/MS

ELOB-associated protein complexes were isolated by FLAG pulldown from BT869 cells expressing FLAG-ELOB or FLAG/HA-GFP (control)(*19*), and by endogenous immunoprecipitation from SU-DIPGXIIIP* cells. Briefly, cells were incubated on ice in swelling buffer (25 mM HEPES pH 7.5, 1.5 mM MgCl₂, 10 mM KCl, 0.05% IGEPAL-CA30) and centrifuged at 300 × g for 10 minutes at 4°C to isolate nuclei. Nuclei were re-suspended in IP buffer (25 mM HEPES, pH 7.5, 150 mM NaCl, 1% Triton X-100) and treated with DNase I (Roche, 04716728001) for 60 minutes on ice, followed by centrifugation at 21,130 × g for 30 minutes at 4°C to remove insoluble material. For FLAG pulldowns, cleared cytosolic and nuclear extracts from BT869 cells were incubated with anti-FLAG M2 resin (Millipore Sigma, A2220) for 3 hours at 4 °C, washed extensively, and eluted with FLAG peptide (100 µg/mL in 10 mM Tris-HCl pH 7.2). Endogenous ELOB complexes were isolated from SU-DIPGXIIIP* extracts by overnight incubation with an ELOB antibody (Abcam, ab45297) or IgG control (Cell Signaling Technology, 2729) at 4°C with rotation, followed by capture with Protein G agarose (Millipore Sigma, 16-266), washing, and elution in RIPA buffer. Histones were isolated by acid extraction and TCA precipitation from control and ELOBKO SU-DIPGXIIIP* and HSJD-DIPG007 cells as previously described (*24*). ELOB protein complexes and histone modification were analyzed by LC–MS/MS on Orbitrap mass spectrometers (ThermoFisher Scientific) at the Taplin Biological Mass Spectrometry Facility at Harvard Medical School.

### Analysis of patient brain tumor transcriptomic and proteomic data

Proteomic and RNA-seq datasets for pediatric high grade glioma (pHGG) were obtained from the PedcBio Portal (https://pedcbioportal.org/) and limited to primary tumors and diagnostic samples.

For patients with multiple samples in the database, mean transcript or protein abundance values were used in downstream analyses. Samples were stratified into expression-defined groups based on the abundance of the protein or transcript interest and correlations with patient survival were determined using Kaplan–Meier analysis in GraphPad Prism. Statistical significance between survival curves was assessed using the log-rank test.

### Quantification and statistical analysis

At least three independent biological replicates or mice are shown in all graphs with individual data points representing individual replicates. Statistical analyses including unpaired student’s t tests, one-way ANOVA, and log-rank tests were performed using GraphPad Prism software. Error bars show S.E.M. with *p<0.05, **p<0.01, ***p<0.001, and ****p<0.0001.

## Supporting information

table s1: CUT&RUN peaks in SU-DIPGXIII following ELOBKO or H3K27MKO

table s2: RNA-seq analysis ELOBKO versus control DIPGXIII, BT245

table s3: PRO-seq analysis ELOBKO versus control DIPGXIII cells

table s4: ELOB IP/MS summary BT869 and SU-DIPGXIIIP* cells

## Acknowledgments

These studies were supported by funding from the National Institutes of Health (5R01N129860), the ChadTough Defeat DIPG Foundation, the Cure Starts Now Foundation, and the Rally Foundation for Childhood Cancer Research and Kids Join the Fight. Dr. Rebecca Murdaugh and Caitlin Bagnetto received training support from the Baylor College of Medicine (BCM) Center for Cell and Gene Therapy (CAGT) training grant (NHLBH, 5T32HL092332). Histone mass spectrometry was performed at Taplin Mass Spectrometry Facility at Harvard Medical School, RNAseq was performed at the genomics core at Tufts University, and PRO-seq analysis was performed through the Nascent Transcriptomics Core at Harvard Medical School.

## Author contributions

Conceptualization: J.N.A. and Y.S. Investigation: A.L.J., R.L.M. A.F.K., C.A.M., B.M.Z., R.U.R. K.Y. and C.L.B. Methodology: A.L.J., R.L.M. A.F.K., C.A.M., B.M.Z., R.U.R. K.Y. and C.L.B. Data Curation: R.L.M. and J.N.A. Validation, formal analysis, software, and visualization: R.L.M., C.A.M. and J.N.A. Project administration: J.N.A. and Y.S. Resources: M.G.F., Y.S., and J.N.A. Writing of the original draft: J.N.A. Review and editing: A.L.J., R.L.M., A.F.K., and Y.S. Funding acquisition and supervision: J.N.A. and Y.S.

## Competing interests

Y.S. is a co-founder of K36 Therapeutics, Alternative Bio (ABio) Inc. and a member of the Scientific Advisory Board of Alternative Bio (ABio) Inc., Epigenica AB and Epic Bio, Inc. Y.S. is also a board member of ABio Inc. and Epigenica AB. Y.S. holds equity in Active Motif, K36 Therapeutics, Epic Bio, Inc., Alternative Bio, Inc. and Epigenica AB. Y.S. serves on the Scientific Advisory Board of the School of Life Sciences, Westlake University and Westlake Laboratory of Life Sciences and Biomedicine, and the Norway Centre for Embryology and Healthy Development. K.A. received research funding from Novartis not related to this work, consults for Syros Pharmaceuticals and Odyssey Therapeutics, and is on the Scientific Advisory Board of CAMP4 Therapeutics.

## Data, code, and materials availability

All data and analysis methods needed to evaluate and reproduce the results in the paper are present in the paper and in the Materials and Methods. Sequencing data generated and analyzed in this work have been deposited in the NCBI Gene Expression Omnibus (GEO) database under the accession numbers: GSE324874: https://www.ncbi.nlm.nih.gov/geo/query/acc.cgi?acc=GSE324874; GSE324875: https://www.ncbi.nlm.nih.gov/geo/query/acc.cgi?acc=GSE3248745; GSE324878: https://www.ncbi.nlm.nih.gov/geo/query/acc.cgi?acc=GSE3248748; H3K4me1, H3K27ac, and H3K27me3 ChIP-seq data from a previous study(*5*) were also used in comparative analyses, which can be found on GEO: GSE110572: https://www.ncbi.nlm.nih.gov/geo/query/acc.cgi?acc=GSE110570 ). All unique plasmid constructs produced in this study are available from the lead contact upon request to J.N.A.

---

**Figure S1.**
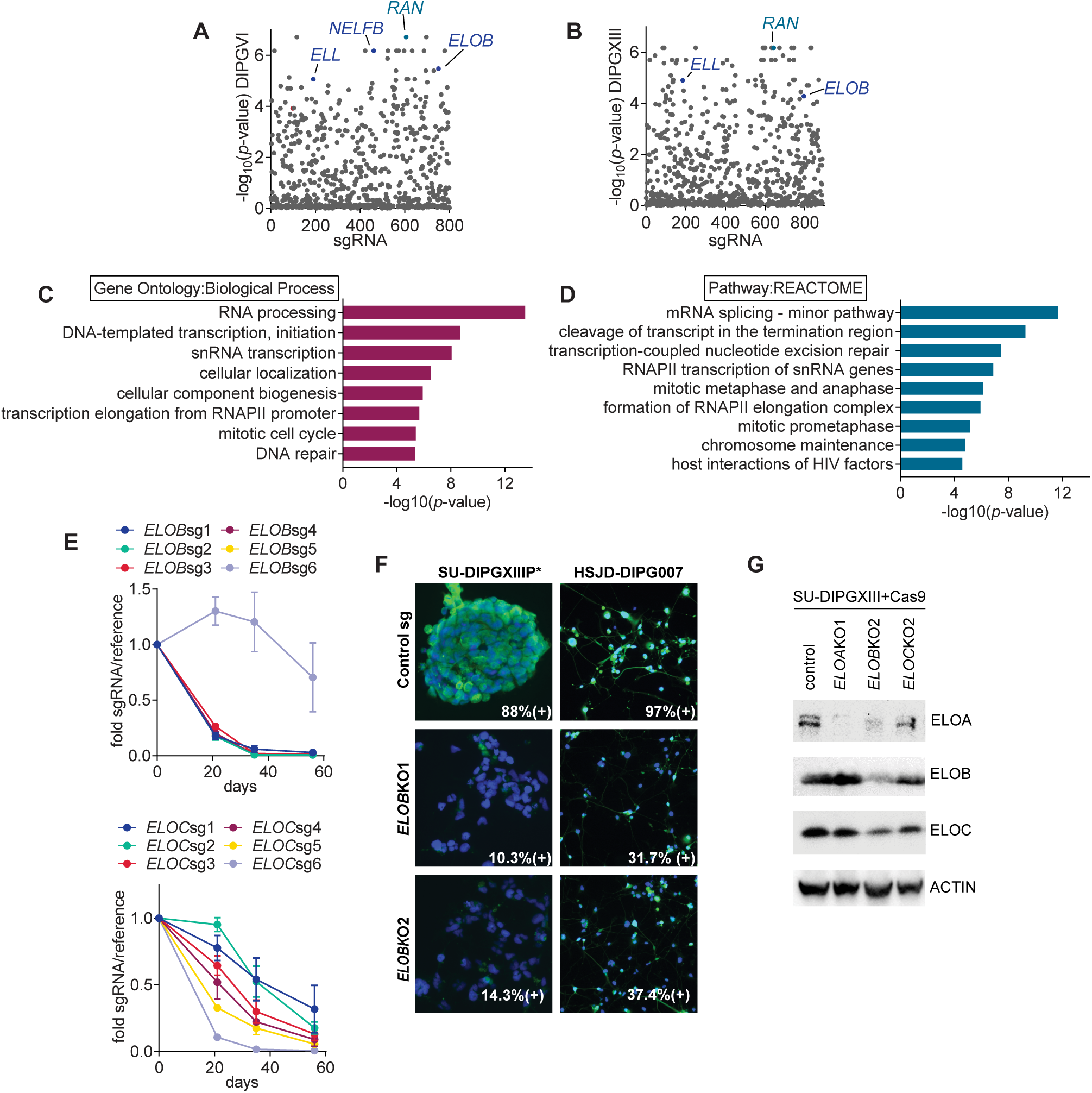
A chromatin-focused CRISPR dropout screen identifies Pol2 transcriptional regulators driving DMG growth. **(A** and **B)** Scatter plots showing –log_10_ (*p*-value) for various genes in the SU-DIPGVI **(A)** and SU-DIPGXIII **(B)** chromatin-focused CRISPR dropout screens calculated using STARS. **(C** and **D)** Enriched gene categories from overrepresentation analysis of screen hits covering the Gene Ontology (Biological Process) **(C)** and REACTOME pathway gene set databases **(D)**. **(E)** Normalized fold change in the expression of 6 independent *ELOB*- (top panel) or *ELOC*-targeting sgRNAs (bottom panel) after 3-8 weeks of SU-DIPGXIII cell outgrowth compared to the 0 week reference sample. **(F)** Representative immunofluorescence images of Cas9 (+) HSJD-DIPG007 and SU-DIPGXIIIP* cells transduced with control and *ELOB* sgRNAs stained to detect endogenous ELOB (green channel) showing ELOB protein loss in a majority of cells. **(G)** Immunoblots from polyclonal SU-DIPGXIII+Cas9 cells transduced with control, *ELOA*, *ELOB*, or *ELOC* sgRNAs confirming partial KO of elongin proteins with a β-actin blot serving as a loading control.

**Figure S2:**
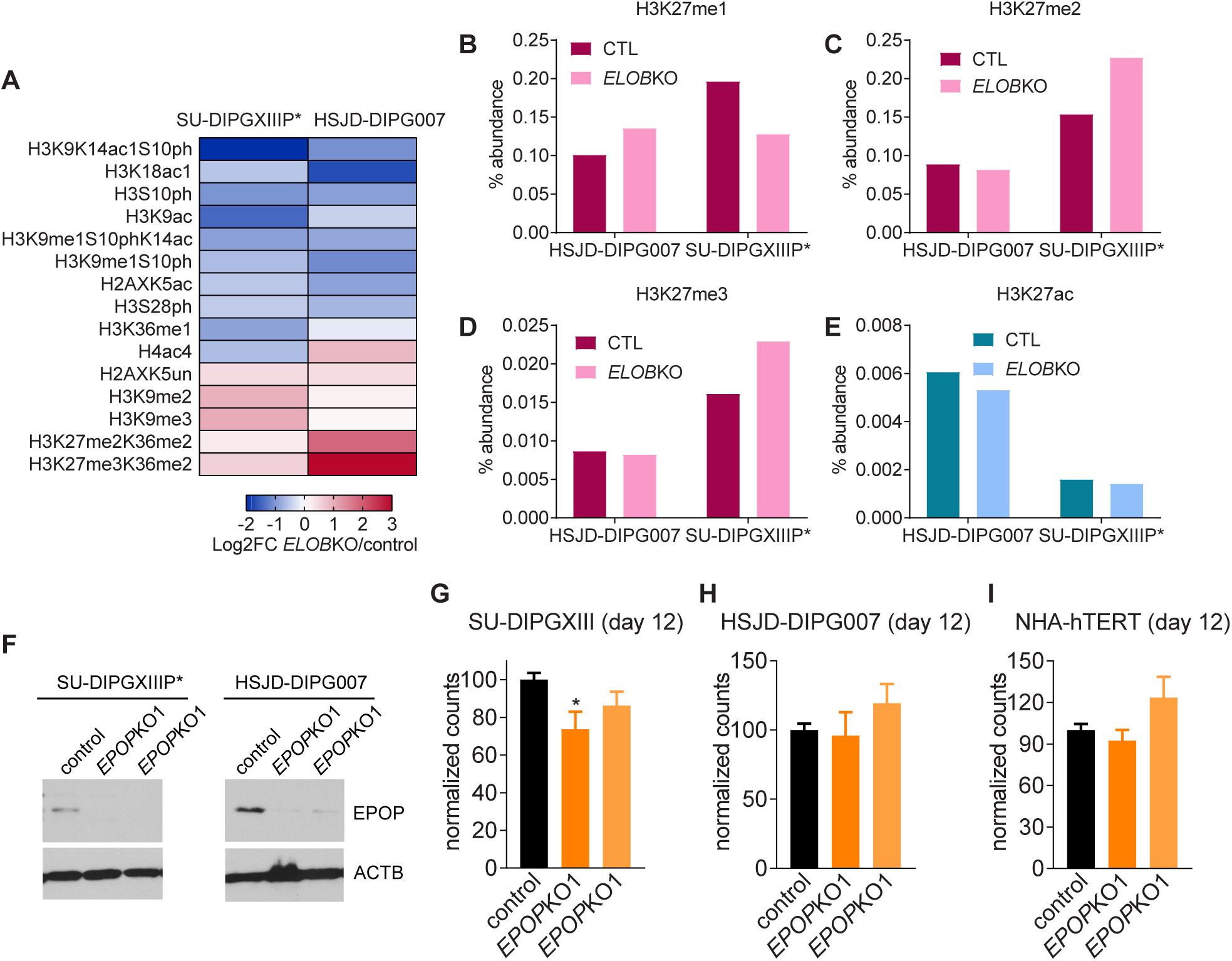
***ELOB*KO alters DMG histone PTMs associated with gene activation and mitosis. (A)** Heatmap showing log_2_ fold change (log_2_FC) for various histone post-translational modifications in *ELOB*KO HSJD-DIPG007 and SU-DIPGXIIIP* cells compared to control sgRNA-transduced cells, revealing reduced H3K9ac, H3K18ac, and H4ac associated with gene activation; reduced H3S10 and H3S28 phosphorylation associated with mitosis; and increased H3K9me2 and H3K9me3 associated with heterochromatin regulation, as determined by tandem mass spectrometry on isolated histones. **(B** to **E)** Percent abundance of PRC2-regulated H3K27me1 **(B)**, -me2 **(C)**, and -me3 **(D)**, or H3K27ac **(E)**, showing no consistent change in H3K27 acetylation or methylation following *ELOB* loss. **(F)** Western blot analysis of lysates from control and *EPOP*KO DMG cells confirming loss of EPOP protein in Cas9 (+) SU-DIPGXIIIP* and HSJD-DIPG007 cells transduced with two independent sgRNAs (*EPOP*KO1 and *EPOP*KO2). **(G** to **I)** Normalized live DMG **(G** and **H)** and immortalized astrocyte (NHA-hTERT, **I**) cell counts 12 days after starting selection for sgRNA transduction to KO *EPOP* suggesting that EPOP-ELOB complexes are not required for optimal DMG cell growth. Error bars show S.E.M. with *p<0.05, **p<0.01, ***p<0.001, and ****p<0.0001.

**Figure S3:**
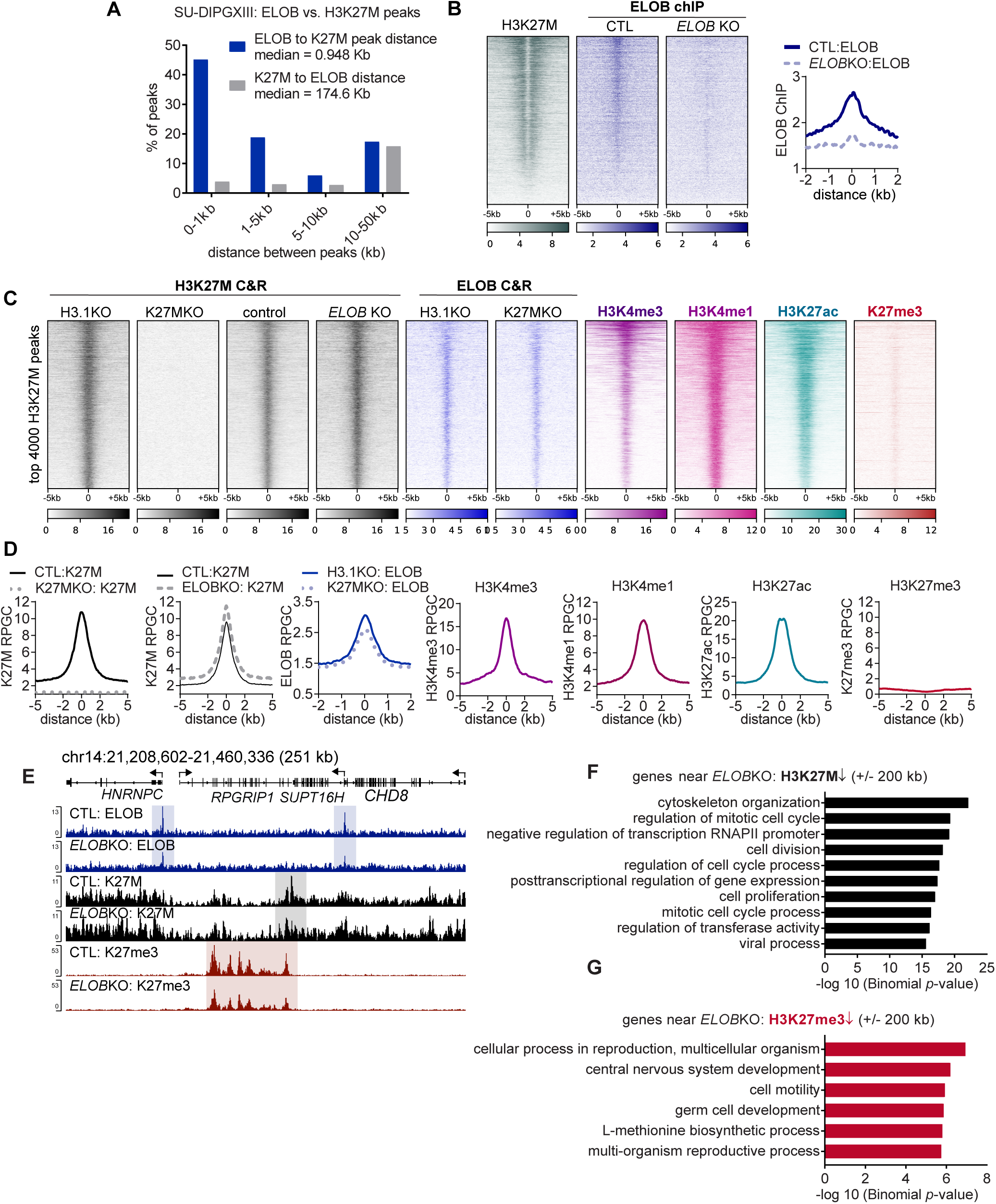
ELOB chromatin binding to active chromatin regions is not dependent on H3K27M. **(A)** Histogram showing the percent of ELOB peaks at various distances (0-50 kb) to the nearest H3K27M peaks (blue bars) or the percent of H3K27M peaks at various distances to ELOB peaks (grey bars), demonstrating proximity between the ELOB peaks and H3K27M-bound regions, consistent with co-localization. **(B)** Heatmap and average profile plot of ELOB ChIP-seq tracks aligned to peaks from ELOB CUT&RUN data (see also Fig. 3A and **B**) further confirming H3K27M co-occupancy at ELOB binding sites. **(C** and **D)** Heatmap **(C)** and average profile plots **(D)** of the top 4,000 H3K27M binding sites identified by CUT&RUN (C&R) showing H3K27M and ELOB chromatin binding profiles in H3.1KO (*HIST1H3B* KO, control) and isogenic H3K27M KO SU-DIPGXIII cells, suggesting that H3K27M is not required for optimal ELOB binding to chromatin. ChIP-seq/CUT&RUN tracks for H3K4me1, H3K4me3, H3K27ac, and H3K27me3 are also included, demonstrating co-enrichment with active chromatin marks and low H3K27me3, as predicted. **(E)** Genome browser snapshot showing ELOB, H3K27M, and H3K27me3 CUT&RUN profiles around the *SUPT16H*, *RPGRIP1*, *HNRNPC*, and *CHD8* gene loci, revealing ELOB peaks (blue box) located in the vicinity of regional losses in H3K27M (gray box) and H3K27me3 binding (red box). **(F** and **G)** GO (biological process) enrichment results shown as –log_10_ binomial *p*-values from an analyses of the single nearest genes within +/- 200 kb of the *ELOB*KO-downregulated H3K27M peaks **(F)** or the *ELOB*KO-downregulated H3K27me3 peaks **(G)**, suggesting that *ELOB* loss alters chromatin state at genes involved in transcriptional regulation, cell division, and central nervous system and germ cell development.

**Figure S4.**
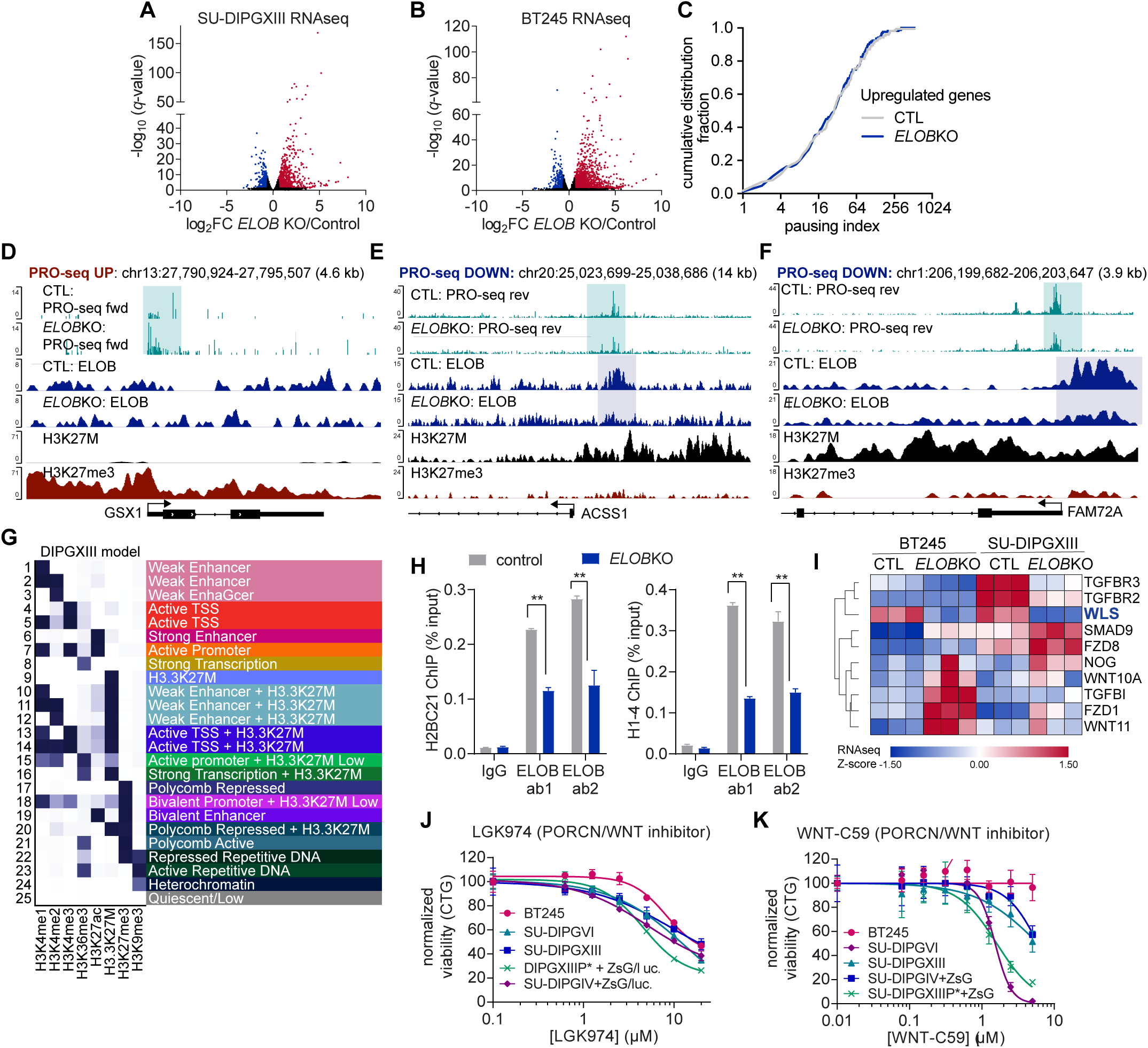
*ELOB*KO alters Pol2 transcriptional activity to regulate DMG gene expression. **(A** and **B)** Volcano plots showing –log_10_ adjusted *p*-value and log_2_ fold change (log_2_FC) from RNA-seq analysis of *ELOB*KO versus control SU-DIPGXIII **(A)** and BT245 cells **(B)**. **(C)** Analysis of the Pol2 pausing index from PRO-seq analysis showing a nonsignificant difference between *ELOB*KO versus control SU-DIPGXIII cells. **(D** to **F)** Genome browser tracks showing increased PRO-seq signal at the H3K27me3-bound *GSX1* gene **(D)** and decreased PRO-seq signal accompanied by ELOB and H3K27M co-occupancy at the *ACSS1* (*Acyl-CoA Synthetase Short Chain Family Member 1,* **E)** and *FAM72A* **(F)** gene loci. **(G)** 25 state ChromHMM model from integrative analysis of chromatin profiling data, PRO-seq, and RNA-seq data in control versus *ELOB*KO DMG cells. **(H)** Summary of ChIP-PCR experiment confirming ELOB binding at *H2BC21* (left) and *H1-4* (right) histone gene loci normalized to input. Chromatin immunoprecipitation was performed with two independent ELOB antibodies (ab1 and ab2) compared to non-specific IgG control. **(I)** Heatmap showing RNA-seq Z-scores of differentially expressed transcripts associated with TGFBβ and WNT signaling in *ELOB*KO versus control DMG cells **(J** and **K).** Normalized viability of various DMG cell lines treated with two structurally distinct inhibitors of WLS/PORCN-dependent WNT secretion (LGK-974, **I** and WNT-C59, **J**) for one week as determined by CellTiter-Glo luciferase assay, suggesting that WNT secretion is required for DMG growth.

**Figure S5.**
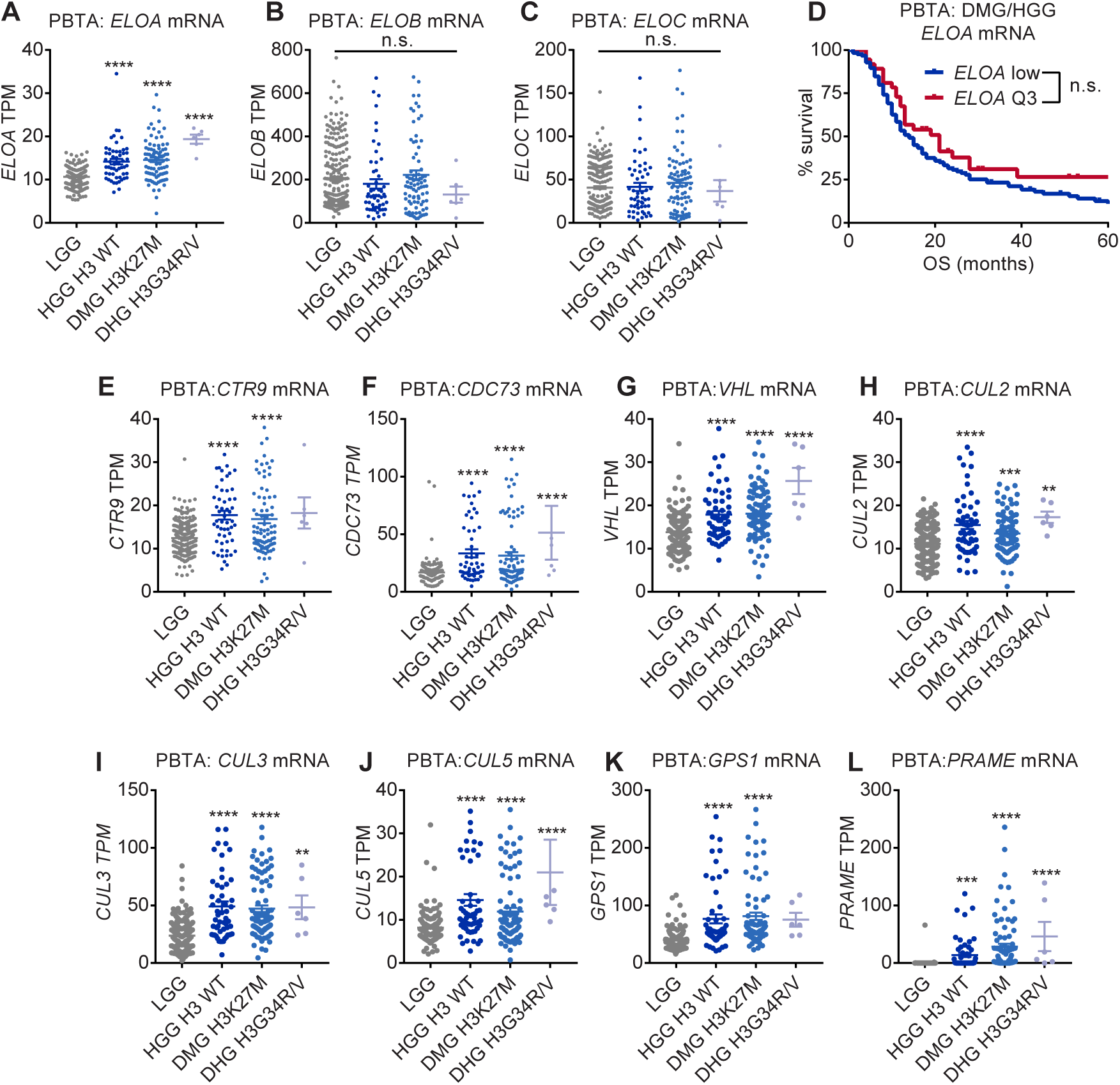
Analysis of ELOB-associated protein expression in pediatric brain tumor clinical samples. **(A** to **C)** mRNA expression of *ELOA* **(A)**, *ELOB* **(B)**, and *ELOC* **(C)** in pediatric high grade glioma (HGG) grouped according to histone H3 mutation status. **(D)** Kaplan-Meier survival analysis showing no significant difference in survival of HGG/DMG patients grouped into high and low *ELOA* expression groups based on publicly available tumor transcriptomics data (PedcBio). **(E** to **L)** Analysis of ELOB-interacting protein expression at the mRNA level in patient LGG and HGG tumors grouped by histone mutation status indicating that multiple ELOB-associated factors are overexpressed in subsets of HGG tumors compared to LGG, but no specific correlation with H3 mutation status.

**table S1: CUT&RUN peaks in SU-DIPGXIII following ELOBKO or H3K27MKO**

**table S2: RNA-seq analysis ELOBKO versus control DIPGXIII, BT245**

**table S3: PRO-seq analysis ELOBKO versus control DIPGXIII cells**

**table S4: ELOB IP/MS summary BT869 and SU-DIPGXIIIP* cells**

## Notes

https://www.ncbi.nlm.nih.gov/geo/query/acc.cgi?acc=GSE324874

https://www.ncbi.nlm.nih.gov/geo/query/acc.cgi?acc=GSE3248745

https://www.ncbi.nlm.nih.gov/geo/query/acc.cgi?acc=GSE3248748

https://www.ncbi.nlm.nih.gov/geo/query/acc.cgi?acc=GSE110570

## References

1. J. Schwartzentruber, A. Korshunov, X.-Y. Liu, D. T. W. Jones, E. Pfaff, K. Jacob, D. Sturm, A. M. Fontebasso, D.-A. K. Quang, M. Tönjes, V. Hovestadt, S. Albrecht, M. Kool, A. Nantel, C. Konermann, A. Lindroth, N. Jäger, T. Rausch, M. Ryzhova, J. O. Korbel, T. Hielscher, P. Hauser, M. Garami, A. Klekner, L. Bognar, M. Ebinger, M. U. Schuhmann, W. Scheurlen, A. Pekrun, M. C. Frühwald, W. Roggendorf, C. Kramm, M. Dürken, J. Atkinson, P. Lepage, A. Montpetit, M. Zakrzewska, K. Zakrzewski, P. P. Liberski, Z. Dong, P. Siegel, A. E. Kulozik, M. Zapatka, A. Guha, D. Malkin, J. Felsberg, G. Reifenberger, A. von Deimling, K. Ichimura, V. P. Collins, H. Witt, T. Milde, O. Witt, C. Zhang, P. Castelo-Branco, P. Lichter, D. Faury, U. Tabori, C. Plass, J. Majewski, S. M. Pfister, N. Jabado, Driver mutations in histone H3.3 and chromatin remodelling genes in paediatric glioblastoma. Nature 482, 226–231 (2012).

2. G. Wu, A. Broniscer, T. A. McEachron, C. Lu, B. S. Paugh, J. Becksfort, C. Qu, L. Ding, R. Huether, M. Parker, J. Zhang, A. Gajjar, M. A. Dyer, C. G. Mullighan, R. J. Gilbertson, E. R. Mardis, R. K. Wilson, J. R. Downing, D. W. Ellison, J. Zhang, S. J. Baker, Somatic histone H3 alterations in pediatric diffuse intrinsic pontine gliomas and non-brainstem glioblastomas. Nat. Genet. 44, 251–253 (2012).

3. A. S. Harutyunyan, B. Krug, H. Chen, S. Papillon-Cavanagh, M. Zeinieh, N. De Jay, S. Deshmukh, C. C. L. Chen, J. Belle, L. G. Mikael, D. M. Marchione, R. Li, H. Nikbakht, B. Hu, G. Cagnone, W. A. Cheung, A. Mohammadnia, D. Bechet, D. Faury, M. K. McConechy, M. Pathania, S. U. Jain, B. Ellezam, A. G. Weil, A. Montpetit, P. Salomoni, T. Pastinen, C. Lu, P. W. Lewis, B. A. Garcia, C. L. Kleinman, N. Jabado, J. Majewski, H3K27M induces defective chromatin spread of PRC2-mediated repressive H3K27me2/me3 and is essential for glioma tumorigenesis. Nat. Commun. 10, 1262 (2019).

4. C. S. Grasso, Y. Tang, N. Truffaux, N. E. Berlow, L. Liu, M.-A. Debily, M. J. Quist, L. E. Davis, E. C. Huang, P. J. Woo, A. Ponnuswami, S. Chen, T. B. Johung, W. Sun, M. Kogiso, Y. Du, L. Qi, Y. Huang, M. Hütt-Cabezas, K. E. Warren, L. Le Dret, P. S. Meltzer, H. Mao, M. Quezado, D. G. van Vuurden, J. Abraham, M. Fouladi, M. N. Svalina, N. Wang, C. Hawkins, J. Nazarian, M. M. Alonso, E. H. Raabe, E. Hulleman, P. T. Spellman, X.-N. Li, C. Keller, R. Pal, J. Grill, M. Monje, Functionally defined therapeutic targets in diffuse intrinsic pontine glioma. Nat. Med. 21, 555–559 (2015).

5. J. N. Anastas, B. M. Zee, J. H. Kalin, M. Kim, R. Guo, S. Alexandrescu, M. A. Blanco, S. Giera, S. M. Gillespie, J. Das, M. Wu, S. Nocco, D. M. Bonal, Q.-D. Nguyen, M. L. Suva, B. E. Bernstein, R. Alani, T. R. Golub, P. A. Cole, M. G. Filbin, Y. Shi, Re-programing Chromatin with a Bifunctional LSD1/HDAC Inhibitor Induces Therapeutic Differentiation in DIPG. Cancer Cell, S1535610819304258 (2019).

6. A. Piunti, R. Hashizume, M. A. Morgan, E. T. Bartom, C. M. Horbinski, S. A. Marshall, E. J. Rendleman, Q. Ma, Y.-H. Takahashi, A. R. Woodfin, A. V. Misharin, N. A. Abshiru, R. R. Lulla, A. M. Saratsis, N. L. Kelleher, C. D. James, A. Shilatifard, Therapeutic targeting of polycomb and BET bromodomain proteins in diffuse intrinsic pontine gliomas. Nat. Med. 23, 493–500 (2017).

7. N. A. Dahl, E. Danis, I. Balakrishnan, D. Wang, A. Pierce, F. M. Walker, A. Gilani, N. J. Serkova, K. Madhavan, S. Fosmire, A. L. Green, N. K. Foreman, S. Venkataraman, R. Vibhakar, Super Elongation Complex as a Targetable Dependency in Diffuse Midline Glioma. Cell Rep. 31, 107485 (2020).

8. S. Nagaraja, N. A. Vitanza, P. J. Woo, K. R. Taylor, F. Liu, L. Zhang, M. Li, W. Meng, A. Ponnuswami, W. Sun, J. Ma, E. Hulleman, T. Swigut, J. Wysocka, Y. Tang, M. Monje, Transcriptional Dependencies in Diffuse Intrinsic Pontine Glioma. Cancer Cell 31, 635–652.e6 (2017).

9. T. E. Miller, B. B. Liau, L. C. Wallace, A. R. Morton, Q. Xie, D. Dixit, D. C. Factor, L. J. Y. Kim, J. J. Morrow, Q. Wu, S. C. Mack, C. G. Hubert, S. M. Gillespie, W. A. Flavahan, T. Hoffmann, R. Thummalapalli, M. T. Hemann, P. J. Paddison, C. M. Horbinski, J. Zuber, P. C. Scacheri, B. E. Bernstein, P. J. Tesar, J. N. Rich, Transcription elongation factors represent in vivo cancer dependencies in glioblastoma. Nature 547, 355–359 (2017).

10. N. A. Dahl, R. Vibhakar, Converging evidence for inhibition of transcriptional control in high-grade gliomas. Neuro-Oncol. 23, 1225–1227 (2021).

11. H. Katagi, N. Takata, Y. Aoi, Y. Zhang, E. J. Rendleman, G. T. Blyth, F. D. Eckerdt, Y. Tomita, T. Sasaki, A. M. Saratsis, A. Kondo, S. Goldman, O. J. Becher, E. Smith, L. Zou, A. Shilatifard, R. Hashizume, Therapeutic targeting of transcriptional elongation in diffuse intrinsic pontine glioma. Neuro-Oncol. 23, 1348–1359 (2021).

12. Z. Luo, C. Lin, A. Shilatifard, The super elongation complex (SEC) family in transcriptional control. Nat. Rev. Mol. Cell Biol. 13, 543–547 (2012).

13. J. N. Bradsher, K. W. Jackson, R. C. Conaway, J. W. Conaway, RNA polymerase II transcription factor SIII. I. Identification, purification, and properties. J. Biol. Chem. 268, 25587–25593 (1993).

14. D. R. Duan, A. Pause, W. H. Burgess, T. Aso, D. Y. T. Chen, K. P. Garrett, R. C. Conaway, J. W. Conaway, W. M. Linehan, R. D. Klausner, Inhibition of Transcription Elongation by the VHL Tumor Suppressor Protein. Science 269, 1402–1406 (1995).

15. T. Aso, W. S. Lane, J. W. Conaway, R. C. Conaway, Elongin (SIII): A Multisubunit Regulator of Elongation by RNA Polymerase II. Science 269, 1439–1443 (1995).

16. A. Shilatifard, W. S. Lane, K. W. Jackson, R. C. Conaway, J. W. Conaway, An RNA Polymerase II Elongation Factor Encoded by the Human ELL Gene. Science 271, 1873–1876 (1996).

17. Y. Yamaguchi, T. Takagi, T. Wada, K. Yano, A. Furuya, S. Sugimoto, J. Hasegawa, H. Handa, NELF, a Multisubunit Complex Containing RD, Cooperates with DSIF to Repress RNA Polymerase II Elongation. Cell 97, 41–51 (1999).

18. Y. Yamaguchi, N. Inukai, T. Narita, T. Wada, H. Handa, Evidence that Negative Elongation Factor Represses Transcription Elongation through Binding to a DRB Sensitivity-Inducing Factor/RNA Polymerase II Complex and RNA. Mol. Cell. Biol. 22, 2918–2927 (2002).

19. R. Liefke, V. Karwacki-Neisius, Y. Shi, EPOP Interacts with Elongin BC and USP7 to Modulate the Chromatin Landscape. Mol. Cell 64, 659–672 (2016).

20. M. B. Ardehali, M. Damle, C. Perea-Resa, M. D. Blower, R. E. Kingston, Elongin A associates with actively transcribed genes and modulates enhancer RNA levels with limited impact on transcription elongation rate in vivo. J. Biol. Chem. 296, 100202 (2021).

21. Y. Wang, L. Hou, M. B. Ardehali, R. E. Kingston, B. D. Dynlacht, Elongin A regulates transcription in vivo through enhanced RNA polymerase processivity. J. Biol. Chem. 296, 100170 (2021).

22. M. Beringer, P. Pisano, V. Di Carlo, E. Blanco, P. Chammas, P. Vizán, A. Gutiérrez, S. Aranda, B. Payer, M. Wierer, L. Di Croce, EPOP Functionally Links Elongin and Polycomb in Pluripotent Stem Cells. Mol. Cell 64, 645–658 (2016).

23. M. Gerber, J. C. Eissenberg, S. Kong, K. Tenney, J. W. Conaway, R. C. Conaway, A. Shilatifard, In Vivo Requirement of the RNA Polymerase II Elongation Factor Elongin A for Proper Gene Expression and Development. Mol. Cell. Biol. 24, 9911–9919 (2004).

24. R. U. Richard, C. Bagnetto, R. L. Murdaugh, B. R. Eberl, A. L. Jiao, A. F. Kebede, B. M. Zee, M. M. Harrington, M. G. Filbin, A. Serin-Harmanci, Y. Shi, J. N. Anastas, SAGA/ATAC complexes sustain aberrant chromatin regulation and promote tumorigenesis in diffuse midline glioma. [Preprint] (2026). 10.64898/2026.01.22.701194.

25. Y. Chen, G. Kokic, C. Dienemann, O. Dybkov, H. Urlaub, P. Cramer, Structure of the transcribing RNA polymerase II–Elongin complex. Nat. Struct. Mol. Biol. 30, 1925–1935 (2023).

26. E. Y. Qin, D. D. Cooper, K. L. Abbott, J. Lennon, S. Nagaraja, A. Mackay, C. Jones, H. Vogel, P. K. Jackson, M. Monje, Neural Precursor-Derived Pleiotrophin Mediates Subventricular Zone Invasion by Glioma. Cell 170, 845–859.e19 (2017).

27. S. Fischer, V. T. Trinh, C. Simon, L. M. Weber, I. Forné, A. Nist, G. Bange, F. Abendroth, T. Stiewe, W. Steinchen, R. Liefke, O. Vázquez, Peptide-mediated inhibition of the transcriptional regulator Elongin BC induces apoptosis in cancer cells. Cell Chem. Biol. 30, 766–779.e11 (2023).

28. F. Mohammad, S. Weissmann, B. Leblanc, D. P. Pandey, J. W. Højfeldt, I. Comet, C. Zheng, J. V. Johansen, N. Rapin, B. T. Porse, A. Tvardovskiy, O. N. Jensen, N. G. Olaciregui, C. Lavarino, M. Suñol, C. de Torres, J. Mora, A. M. Carcaboso, K. Helin, EZH2 is a potential therapeutic target for H3K27M-mutant pediatric gliomas. Nat. Med. 23, 483–492 (2017).

29. M. P. Meers, T. D. Bryson, J. G. Henikoff, S. Henikoff, Improved CUT&RUN chromatin profiling tools. eLife 8 (2019).

30. A. S. Harutyunyan, H. Chen, T. Lu, C. Horth, H. Nikbakht, B. Krug, C. Russo, E. Bareke, D. M. Marchione, M. Coradin, B. A. Garcia, N. Jabado, J. Majewski, H3K27M in Gliomas Causes a One-Step Decrease in H3K27 Methylation and Reduced Spreading within the Constraints of H3K36 Methylation. Cell Rep. 33, 108390 (2020).

31. T. Kouzarides, Chromatin modifications and their function. Cell 128, 693–705 (2007).

32. H. Huang, B. R. Sabari, B. A. Garcia, C. D. Allis, Y. Zhao, SnapShot: Histone Modifications. Cell 159, 458–458.e1 (2014).

33. W. Alafate, G. Lv, J. Zheng, H. Cai, W. Wu, Y. Yang, S. Du, D. Zhou, P. Wang, Targeting ARNT attenuates chemoresistance through destabilizing p38α-MAPK signaling in glioblastoma. Cell Death Dis. 15, 366 (2024).

34. S. K. Sgaier, Z. Lao, M. P. Villanueva, F. Berenshteyn, D. Stephen, R. K. Turnbull, A. L. Joyner, Genetic subdivision of the tectum and cerebellum into functionally related regions based on differential sensitivity to engrailed proteins. Development 134, 2325–2335 (2007).

35. Y. Wu, M. Fletcher, Z. Gu, Q. Wang, B. Costa, A. Bertoni, K.-H. Man, M. Schlotter, J. Felsberg, J. Mangei, M. Barbus, A.-C. Gaupel, W. Wang, T. Weiss, R. Eils, M. Weller, H. Liu, G. Reifenberger, A. Korshunov, P. Angel, P. Lichter, C. Herrmann, B. Radlwimmer, Glioblastoma epigenome profiling identifies SOX10 as a master regulator of molecular tumour subtype. Nat. Commun. 11, 6434 (2020).

36. A. Piunti, R. Hashizume, M. A. Morgan, E. T. Bartom, C. M. Horbinski, S. A. Marshall, E. J. Rendleman, Q. Ma, Y.-H. Takahashi, A. R. Woodfin, A. V. Misharin, N. A. Abshiru, R. R. Lulla, A. M. Saratsis, N. L. Kelleher, C. D. James, A. Shilatifard, Therapeutic targeting of polycomb and BET bromodomain proteins in diffuse intrinsic pontine gliomas. Nat. Med. 23, 493–500 (2017).

37. J. Ernst, M. Kellis, ChromHMM: automating chromatin-state discovery and characterization. Nat. Methods 9, 215–216 (2012).

38. J. D. Larson, L. H. Kasper, B. S. Paugh, H. Jin, G. Wu, C.-H. Kwon, Y. Fan, T. I. Shaw, A. B. Silveira, C. Qu, R. Xu, X. Zhu, J. Zhang, H. R. Russell, J. L. Peters, D. Finkelstein, B. Xu, T. Lin, C. L. Tinkle, Z. Patay, A. Onar-Thomas, S. B. Pounds, P. J. McKinnon, D. W. Ellison, J. Zhang, S. J. Baker, Histone H3.3 K27M Accelerates Spontaneous Brainstem Glioma and Drives Restricted Changes in Bivalent Gene Expression. Cancer Cell 35, 140–155.e7 (2019).

39. B. E. Bernstein, T. S. Mikkelsen, X. Xie, M. Kamal, D. J. Huebert, J. Cuff, B. Fry, A. Meissner, M. Wernig, K. Plath, R. Jaenisch, A. Wagschal, R. Feil, S. L. Schreiber, E. S. Lander, A Bivalent Chromatin Structure Marks Key Developmental Genes in Embryonic Stem Cells. Cell 125, 315–326 (2006).

40. J. Liu, S. Pan, M. H. Hsieh, N. Ng, F. Sun, T. Wang, S. Kasibhatla, A. G. Schuller, A. G. Li, D. Cheng, J. Li, C. Tompkins, A. Pferdekamper, A. Steffy, J. Cheng, C. Kowal, V. Phung, G. Guo, Y. Wang, M. P. Graham, S. Flynn, J. C. Brenner, C. Li, M. C. Villarroel, P. G. Schultz, X. Wu, P. McNamara, W. R. Sellers, L. Petruzzelli, A. L. Boral, H. M. Seidel, M. E. McLaughlin, J. Che, T. E. Carey, G. Vanasse, J. L. Harris, Targeting Wnt-driven cancer through the inhibition of Porcupine by LGK974. Proc. Natl. Acad. Sci. 110, 20224–20229 (2013).

41. K. D. Proffitt, B. Madan, Z. Ke, V. Pendharkar, L. Ding, M. A. Lee, R. N. Hannoush, D. M. Virshup, Pharmacological inhibition of the Wnt acyltransferase PORCN prevents growth of WNT-driven mammary cancer. Cancer Res. 73, 502–507 (2013).

42. Y. Zhang, F. Chen, L. A. Donehower, M. E. Scheurer, C. J. Creighton, A pediatric brain tumor atlas of genes deregulated by somatic genomic rearrangement. Nat. Commun. 12, 937 (2021).

43. H. Ijaz, M. Koptyra, K. S. Gaonkar, J. L. Rokita, V. P. Baubet, L. Tauhid, Y. Zhu, M. Brown, G. Lopez, B. Zhang, S. J. Diskin, Z. Vaksman, Children’s Brain Tumor Tissue Consortium, J. L. Mason, E. Appert, J. Lilly, R. Lulla, T. De Raedt, A. P. Heath, A. Felmeister, P. Raman, J. Nazarian, M. R. Santi, P. B. Storm, A. Resnick, A. J. Waanders, K. A. Cole, Pediatric high-grade glioma resources from the Children’s Brain Tumor Tissue Consortium. Neuro-Oncol. 22, 163–165 (2020).

44. A. Ranjan, Y. Pang, M. Butler, M. Merchant, O. Kim, G. Yu, Y.-T. Su, M. R. Gilbert, D. Levens, J. Wu, Targeting CDK9 for the Treatment of Glioblastoma. Cancers 13, 3039 (2021).

45. J. Wu, Y. Yuan, D. A. Long Priel, D. Fink, C. J. Peer, T. M. Sissung, Y.-T. Su, Y. Pang, G. Yu, M. K. Butler, T. R. Mendoza, E. Vera, S. Ahmad, C. Bryla, M. Lindsley, E. Grajkowska, K. Mentges, L. Boris, R. Antony, N. Garren, C. Siegel, N. Lollo, C. Cordova, O. Aboud, B. J. Theeler, E. M. Burton, M. Penas-Prado, H. Leeper, J. Gonzales, T. S. Armstrong, K. R. Calvo, W. D. Figg, D. B. Kuhns, J. I. Gallin, M. R. Gilbert, Phase I Study of Zotiraciclib in Combination with Temozolomide for Patients with Recurrent High-grade Astrocytomas. Clin. Cancer Res. 27, 3298–3306 (2021).

46. S. Guo, Y. Yamaguchi, S. Schilbach, T. Wada, J. Lee, A. Goddard, D. French, H. Handa, A. Rosenthal, A regulator of transcriptional elongation controls vertebrate neuronal development. Nature 408, 366–369 (2000).

47. T. Yasukawa, S. Bhatt, T. Takeuchi, J. Kawauchi, H. Takahashi, A. Tsutsui, T. Muraoka, M. Inoue, M. Tsuda, S. Kitajima, R. C. Conaway, J. W. Conaway, P. A. Trainor, T. Aso, Transcriptional elongation factor elongin A regulates retinoic acid-induced gene expression during neuronal differentiation. Cell Rep. 2, 1129–1136 (2012).

48. J. Li, X. Xu, M. Tiwari, Y. Chen, M. Fuller, V. Bansal, P. Tamayo, S. Das, P. Ghosh, G. L. Sen, SPT6 promotes epidermal differentiation and blockade of an intestinal-like phenotype through control of transcriptional elongation. Nat. Commun. 12, 784 (2021).

49. X. Bai, J. Kim, Z. Yang, M. J. Jurynec, T. E. Akie, J. Lee, J. LeBlanc, A. Sessa, H. Jiang, A. DiBiase, Y. Zhou, D. J. Grunwald, S. Lin, A. B. Cantor, S. H. Orkin, L. I. Zon, TIF1gamma controls erythroid cell fate by regulating transcription elongation. Cell 142, 133–143 (2010).

50. X. Liu, W. L. Kraus, X. Bai, Ready, pause, go: regulation of RNA polymerase II pausing and release by cellular signaling pathways. Trends Biochem. Sci. 40, 516–525 (2015).

51. Y. He, S. Sato, C. Tomomori-Sato, S. Chen, Z. H. Goode, J. W. Conaway, R. C. Conaway, Elongin functions as a loading factor for Mediator at ATF6α-regulated ER stress response genes. Proc. Natl. Acad. Sci. 118, e2108751118 (2021).

52. M. J. Guertin, S. J. Petesch, K. L. Zobeck, I. M. Min, J. T. Lis, Drosophila Heat Shock System as a General Model to Investigate Transcriptional Regulation. Cold Spring Harb. Symp. Quant. Biol. 75, 1–9 (2010).

53. S. Henikoff, Y. Zheng, R. M. Paranal, Y. Xu, J. E. Greene, J. G. Henikoff, Z. R. Russell, F. Szulzewsky, H. N. Thirimanne, S. Kugel, E. C. Holland, K. Ahmad, RNA polymerase II at histone genes predicts outcome in human cancer. Science 387, 737–743 (2025).

54. F. Robert, C. Jeronimo, Transcription-coupled nucleosome assembly. Trends Biochem. Sci. 48, 978–992 (2023).

55. R. Franklin, B. Zhang, J. Frazier, M. Chen, B. T. Do, S. Padayao, K. Wu, M. G. Vander Heiden, C. R. Vakoc, J.-S. Roe, M. Ninova, J. Murn, D. B. Sykes, S. Cheloufi, Histone chaperones coupled to DNA replication and transcription control divergent chromatin elements to maintain cell fate. Genes Dev. 39, 652–675 (2025).

56. B. Chen, R. Dronamraju, W. R. Smith-Kinnaman, S. A. Peck Justice, A. J. Hepperla, H. K. MacAlpine, J. M. Simon, A. L. Mosley, D. M. MacAlpine, B. D. Strahl, Spt6-Spn1 interaction is required for RNA polymerase II association and precise nucleosome positioning along transcribed genes. J. Biol. Chem. 301, 108436 (2025).

57. S. Arfè, T. Karagyozova, A. Forest, D. Bingham, H. Hmidan, D. Mazaud, M. Garnier, P. Le Baccon, E. Meshorer, J.-P. Quivy, G. Almouzni, H3.3 deposition counteracts the replication-dependent enrichment of H3.1 at chromocenters in embryonic stem cells. Nat. Commun. 16, 5138 (2025).

58. S. Mendiratta, A. Gatto, G. Almouzni, Histone supply: Multitiered regulation ensures chromatin dynamics throughout the cell cycle. J. Cell Biol. 218, 39–54 (2019).

59. A. D. Goldberg, L. A. Banaszynski, K.-M. Noh, P. W. Lewis, S. J. Elsaesser, S. Stadler, S. Dewell, M. Law, X. Guo, X. Li, D. Wen, A. Chapgier, R. C. DeKelver, J. C. Miller, Y.-L. Lee, E. A. Boydston, M. C. Holmes, P. D. Gregory, J. M. Greally, S. Rafii, C. Yang, P. J. Scambler, D. Garrick, R. J. Gibbons, D. R. Higgs, I. M. Cristea, F. D. Urnov, D. Zheng, C. D. Allis, Distinct factors control histone variant H3.3 localization at specific genomic regions. Cell 140, 678–691 (2010).

60. G. Li, X. Lin, X. Wang, L. Cai, J. Liu, Y. Zhu, Z. Fu, Enhancing radiosensitivity in triple-negative breast cancer through targeting ELOB. Breast Cancer 31, 426–439 (2024).

61. L. Tian, L. Gong, C. Hao, Y. Feng, S. Yao, B. Fei, X. Wang, Z. Huang, ELOA promotes tumor growth and metastasis by activating RBP1 in gastric cancer. Cancer Med. 12, 18946–18959 (2023).

62. C. A. Mimoso, S. R. Goldman, PRO-seq: Precise Mapping of Engaged RNA Pol II at Single-Nucleotide Resolution. Curr. Protoc. 3, e961 (2023).

